# Combining evolution and machine learning-guided pathway optimization to engineer a novel methylsuccinate module for synthetic C1 metabolism *in vivo*

**DOI:** 10.64898/2026.01.16.699985

**Authors:** Helena Schulz-Mirbach, Vittorio Rainaldi, Nitin Bohra, Katsutaka Suzuki, Titouan Danet, Huda Kasim, Ari Satanowski, Hai He, Elena Rossini, Seung Hwan Lee, Melanie Klose, Jörg Kahnt, Timo Glatter, Peter Claus, Nicole Paczia, Beau Dronsella, Shanshan Luo, Nico J. Claassens, Tobias J. Erb

## Abstract

*De novo* metabolic pathways open possibilities for sustainable biotransformations in microbes. However, the *in vivo*-implementation of such new-to-nature pathways is highly challenging and heavily relies on adaptive laboratory evolution (ALE) of the host’s native metabolic network. Here, we assess how much this need for host-centric ALE can be overcome and/or complemented through the informed design of the newly introduced pathway. Exemplifying for a synthetic CO_2_-fixation module via methyl-succinate, we established methylsuccinate-dependent growth of *Escherichia coli* over six months by ALE of *E. coli*’s native metabolism. In parallel, we developed a machine-learning guided workflow (MEVIS) for the automated engineering of the synthetic pathway, resulting in methylsuccinate-dependent growth within three weeks. Critically, performing MEVIS in the background of the ALE-evolved strain is necessary to further approach wild-type like growth, demonstrating how ALE in combination with machine-learning-guided lab automation holds great potential to accelerate and improve design-build-test-learn cycles in contemporary metabolic engineering.

## Introduction

Synthetic metabolism aims at developing orthogonal, i.e. “new-to-nature”, pathways that are more efficient than those evolved by nature. One prominent example are synthetic CO_2_-fixation pathways. These pathways capture the greenhouse gas CO_2_ and were specifically designed to overcome the limitations of natural CO_2_-capturing solutions, such as the Calvin-Benson-Bassham cycle of photosynthesis, which is the dominant CO_2_-fixation process on Earth, but suffers from low turnover rates and high energetic costs^1,2^.

Several synthetic CO_2_-fixation pathways with varying degrees of orthogonality have been proposed ^3–6^, and first successful transfers of simpler designs – or parts thereof, so-called modules – into model microbes were reported^7–9^. ^1,10,11^ Current strategies for the *in vivo* implementation of synthetic pathways are based on introducing defined auxotrophies, which force the host cell to rely on the synthetic pathway – or a pathway module of interest – for growth and survival. In such growth-coupled selections, the growth phenotype is a readout for flux through the synthetic pathway of interest, where growth rate ideally directly correlates with pathway activity ^12,13^.

One example is the implementation of the reductive glycine pathway into the chemolithotroph *Cupriavidus necator*, which increased CO_2_-dependent biomass formation by ∼20% compared to the natural Calvin cycle^1^. Yet, even the transfer of a relatively “simple” design like the reductive glycine pathway, which was constructed around natural glycine/serine transformations, required extensive Design-Build-Test-Learn (DBTL) cycles for successful implementation. In case of the reductive glycine pathway, which consists of “only” four additional enzymes and directly feeds into central metabolism, full re-wiring took more than five years ^10,11,14–19^.

In contrast to the above example, more orthogonal designs for synthetic CO_2_ fixation did not yet support engineered autotrophy, thus far. Critically, many of the most promising pathway designs rely on the activity of crotonyl-CoA carboxylase/reductase (Ccr). This enzyme belongs to the family of enoyl-CoA carboxylase/reductases and is one of the fastest CO_2_-converting biocatalysts described to date^3,5^. Ccr is at the core of various synthetic designs, including the CETCH^3^, HOPAC^20^, rGS-MCG^4^ and THETA^9^ cycles (Figure 1). While various Ccr-based pathways have been successfully reconstructed in cell-free systems^4,9,20,20^, their transfer into living cells has still not been achieved, highlighting our limitations to realize truly “synthetic” metabolism in living cells.

**Figure 1:**
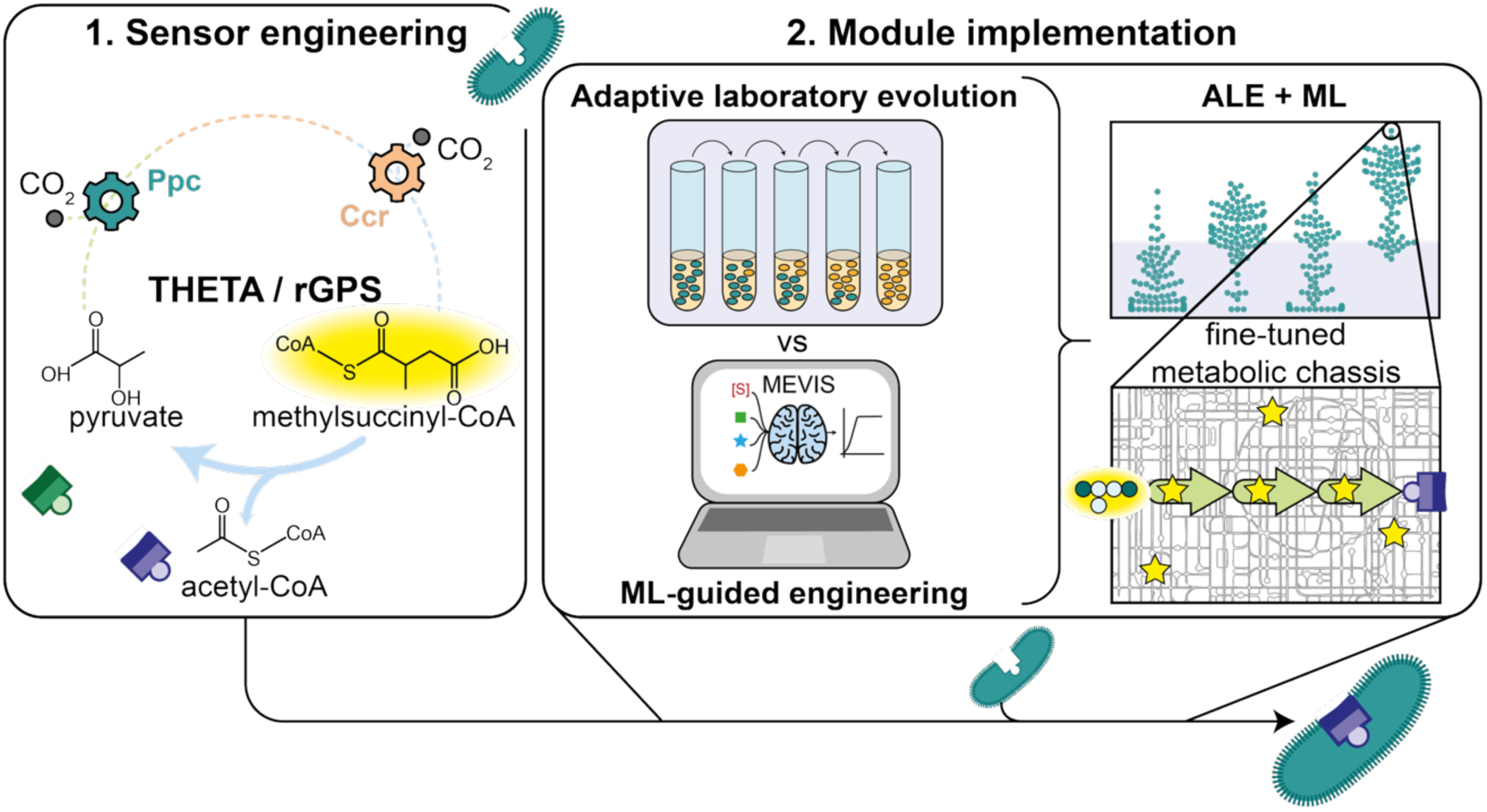
Ccr allows synthetic CO_2_ fixation in the THETA and rGPS cycles. To couple cell growth to methylsuccinyl-CoA as key metabolite downstream of Ccr, we first created sensor strains for the respective module outputs pyruvate and acetyl-CoA. Methylsuccinate dependency of an acetate sensor strain was achieved individually via adaptive laboratory evolution (ALE) and the MEVIS workflow for machine-learning guided (ML) pathway engineering, which were ultimately combined for the creation of a metabolically optimized methylsuccinate sensor. Colored gear wheels indicate carboxylases.

One main challenge in these efforts is the fact that highly orthogonal pathways have little overlap with natural metabolism. Critically, a high degree of pathway orthogonality requires more elaborate metabolic rewiring to achieve such growth-coupling^8,21^. In case of Ccr, multiple of the metabolites downstream are entirely foreign to *E. coli*. Connecting such a metabolic module with the endogenous metabolism of the host requires functionalizing long reaction cascades (Figure 1)^13,22^, which involves tedious DBTL cycles and often requires experimental evolution to functionally connect the synthetic and natural metabolic networks. In case of Ccr-based designs, a first success in a 2-oxoglutarate auxotrophic strain was recently reported. Yet, achieving growth via a Ccr-based module required multiple steps of adaptive laboratory evolution (ALE), with the underlying mechanisms finally allowing growth remaining unclear^8^.

Beyond this example, ALE has proven essential for the implementation of most synthetic pathways^10,11,14,15,23^. Thus, current engineering strategies in synthetic metabolism are still empirical and often use “black box” approaches that depend on spontaneous mutations for pathway-dependent growth^14,15,23^. Critically, to what extent host cells need to adopt their native metabolic and regulatory networks to accept a synthetic pathway—and, *vice versa*, to which extent newly introduced networks need to be fine-tuned to integrate into native metabolism is less understood, let alone considered, during implementation.

In this study, we aimed at overcoming these fundamental shortcomings in synthetic pathway engineering by systematically approaching implementation from both, the side of the host, as well as the synthetic metabolic module. To that end, we first developed an array of selection strains suitable for the growth-coupled implementation of different Ccr-based pathway modules. Exemplifying for the methylsuccinate module of the THETA cycle, we show how evolutionary adaptations of the host can accommodate a newly introduced synthetic module and pave the way for growth-coupling over the course of six months. In parallel to these host-centric efforts, we developed a machine learning-guided workflow for the semi-automated (fine-)tuning of synthetic modules (MEVIS). Applying MEVIS allowed us to achieve growth-coupling through optimization of the methylsuccinate module in the background of the selection strain within (just) four weeks. Importantly, we also show that while ALE of the host as well as MEVIS-guided optimization of the synthetic pathway are each sufficient to yield robust basic growth through the synthetic pathway module (growth rate of ∼0.1 h^-1^), their combination is synergistic and necessary to further improve methylsuccinate-dependent growth towards wild-type like growth (growth rate of ∼0.2 h^-1^).

In summary, our work provides a first systematic and comparative effort towards engineering of orthogonal metabolic routes into host cells, demonstrating the relevance of combining host-centric approaches (experimental evolution) and module-centric approaches (machine learning-guided optimization) to achieve functional implementation. For the latter, we introduce MEVIS, as a convenient and rapid workflow to search the expression space of synthetic pathway modules, which is not limited to the proof-of-principle application in this study, but has the potential to significantly aid and accelerate metabolic engineering beyond synthetic CO_2_-fixation in the future.

## Results

### Sensor strains selecting for Ccr-dependent fixation of CO_2_ into methylsuccinyl-CoA

First, we sought to construct selection strains that would allow for the implementation of the Ccr-module of the THETA and rGPS cycles (Figure 1). This required interfacing methylsuccinyl-CoA, the product of the Ccr reaction, with native *E. coli* metabolism to achieve growth dependency. In the THETA and rGPS cycles, the first *E. coli* metabolites downstream of Ccr are acetyl-CoA and pyruvate (THETA), which are obtained through cleavage of citramalyl-CoA (Figure 1)^6,9^. For acetyl-CoA-based selection, we aimed at utilizing a previously published acetyl-CoA auxotrophic strain (SL2)^9^. The SL2 strain requires acetyl-CoA for the biosynthesis of leucine and fatty acids, as well as for synthesizing reducing equivalents and 2-oxoglutarate via acetyl-CoA oxidation in the TCA cycle (Figure 2, Extended Data Figure 1A).

**Figure 2:**
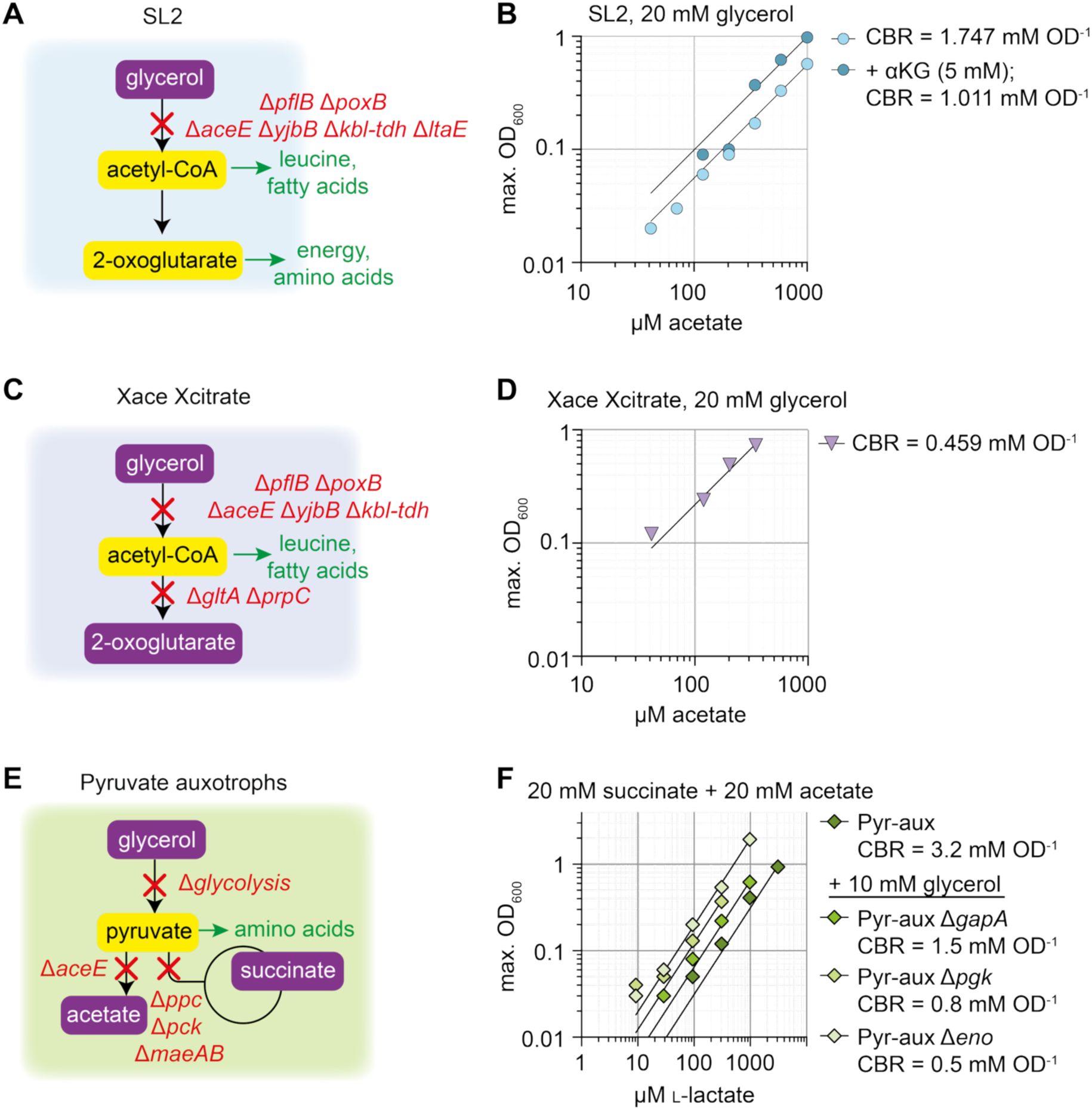
Sensor strains created and characterized to engineer Ccr dependency. **A)** The SL2 strain was previously published by Luo et al. ^9^ and relies on acetyl-CoA for the synthesis of leucine, fatty acids, and TCA cycle intermediates. **B)** Sensitivity characterization of the SL2 strain with and without 2-oxoglutarate supplementation. CBR indicates the compound biomass ratio, in this case how much acetyl-CoA (mM) is required to achieve an optical density of 1. **C)** The Xace Xcitrate strain does not incorporate acetyl-CoA into the TCA cycle anymore due to the deletions of *gltA* and *prpC* and therefore relies on 2-oxoglutarate supplementation. **D)** The Xace Xcitrate strain is more sensitive than the SL2 strain. 20 mM glycerol and 5 mM 2-oxoglutarate were supplied for strain growth. **E)** Pyruvate auxotrophic strains were created by isolating pyruvate from glycolysis and the TCA cycle. **F)** Four created Pyruvate auxotrophs allow sensing lactate in concentrations between 50 µM and 10 mM.

We first characterized acetate sensitivity and substrate co-supplementation of the SL2 strain (Figure 2A, B), and verified its stability over prolonged periods of incubation (250 hours, Extended Data Figure 1B). Upon 2-oxoglutarate supplementation, we observed a 40 % increased sensitivity of the strain for acetyl-CoA (indicated by the compound biomass ratio (CBR)^13^, Figure 2B, Extended Data Figure 1B). Thus, we reasoned that supplementing 2-oxoglutarate in combination with deletion of citrate synthase (Δ*gltA* Δ*prpC*) would further reduce acetyl-CoA funneling into the TCA cycle. Indeed, the resulting strain (named Xace Xcitrate after the disruptions of the acetyl-CoA and citrate nodes) carrying all deletions of the SL2 strain as well as Δ*gltA* Δ*prpC* deletions (Extended Data Figure 1C) grew to 2-fold higher optical densities with low acetate concentrations (Figure 2C, D, Extended Data Figure 1D) and was significantly more sensitive compared to the SL2 strain.

Parallel to above acetyl-CoA auxotrophs, we created pyruvate auxotrophic strains that could select for reactions of the THETA and rGPS modules downstream of Ccr. These strains were designed to exhibit different degrees of pyruvate dependency, for which we metabolically isolated pyruvate, the TCA cycle, and different parts of glycolysis (Figure 2E). Based on the point of intervention, the number of metabolites, the percentage of biomass, and thus selection pressure, increases, which allowed us to tune the selective demand of the different strains (Extended Data Figure 2A). The final four strains: Pyr-aux. (expected to select for 52 % biomass^24,25^), Pyr-aux. Δ*gapA* <u>and</u> Pyr-aux. Δ*pgk* (expected to select for 34 % of biomass^24,25^), and Pyr-aux. Δ*eno* (expected to select for 24 % of biomass^24,25^), cover a concentration range of two orders of magnitude for pyruvate (50 µM to 3 mM, tested by feeding lactate, which is converted to pyruvate by *E. coli*; Figure 2F, Extended Data Figure 2B, C). We additionally tested the effect of co-supplementing pyruvate-derived amino acids on sensitivity of these strains. Supplementing the most sensitive Pyr-aux. Δ*eno* strain with a mixture of lysine, isoleucine, leucine, and valine (K, I, L, V), which are derived from pyruvate but cannot be converted back to it, extended sensitivity of the Pyr-aux. Δ*eno* strain by another twofold (Extended Data Figure 2D, E).

### Developing a CoA-independent methylsuccinate assimilation route

Having established the various auxotrophic strains, we next sought to test growth via the Ccr-THETA module in these strains. Expression of a CoA ligase (Q9HWI3), Ccr, ethylmalonyl-CoA mutase (Ecm), ethylmalonyl-CoA isomerase (Epi), a CoA transferase (Ict-Pa), mesaconyl-C4-CoA hydratase (Meh), mesaconyl-C1-CoA-C4-CoA transferase (Mct), and a citramalyl-CoA lyase (Ccl) did not result in growth with crotonate as carbon source, neither in the acetyl-CoA auxotrophic SL2 strain, nor in the Xace Xcitrate derivative, which is in line with earlier observations^9^.

Similarly, we also did not observe growth on crotonate for the highly sensitive Pyr-aux Δ*eno strain*, even when supplementing amino acids KILV to further decrease the selective demand in this strain by another twofold (Extended Data Figure 3A, B). Critically, when feeding methylsuccinate and mesaconate that are directly down-stream metabolites of the Ccr reaction, and that are activated into the respective CoA esters via Ict-Pa *in vivo*^26^, we did not observe growth of the Pyr-aux. Δ*pgk* strain (Extended Data Figure 4C). These observations pointed to a more general issue of establishing methylsuccinyl- and mesaconyl-CoA transformations in *E. coli*. Note that this finding is in line with previous efforts of implementing the THETA cycle into *E. coli*, which indicated that the inherent instability of methylsuccinyl-CoA and the presence of native *E. coli* thioesterases, such as YciA and TesB, strongly affect CoA ester of the methylsuccinyl-CoA conversion module^9^. Thus, we decided to develop an acid-based instead of a CoA-ester based route for the downstream transformation of methyl-succinate (Supplementary Figure 1A).

For the CoA-based route, to convert methylsuccinyl-CoA to mesaconyl-CoA, we employed a methylsuccinyl-CoA oxidase (Supplementary Figure 1A). Switching to a acid-based route, required us to identify a corresponding methylsuccinate oxidase or dehydrogenase. In search for suitable candidates, we assessed recent efforts of engineering methylsuccinate dependency: Luo et al. recently found that native succinate dehydrogenase (SdhA) of *E. coli* has basal activity with methylsuccinate^9,20^ (Figure 3A). Rainaldi et al. engineered the reductive methylaspartate cycle into *E. coli*, where they identified a F119L variant of SdhA that emerged during evolution towards methylsuccinate dependency^8^. Another instance of methylsuccinate dependency was recently reported by Hwan Lee et al., who evolved strain IDM-P for growth on methylsuccinate^26^. Upon sequencing the evolved isolate, we found that the evolved isolate carried the same SdhA F119L mutation that was found by Rainaldi et al. (Supplementary Dataset 1). When testing the activity of the enzyme *in vitro*, we observed an ∼40% increased apparent turnover and a ∼40% lower *K_M_* value for methylsuccinate of the F119L SdhA mutant variant with methylsuccinate, resulting in about two-fold increased catalytic efficiency compared to the wild-type enzyme (Figure 3A-C, Supplementary Figure 1B). Thus, we decided to use SdhA F119L as methyl-succinate dehydrogenase in our effort to establish the Ccr-THETA module.

**Figure 3:**
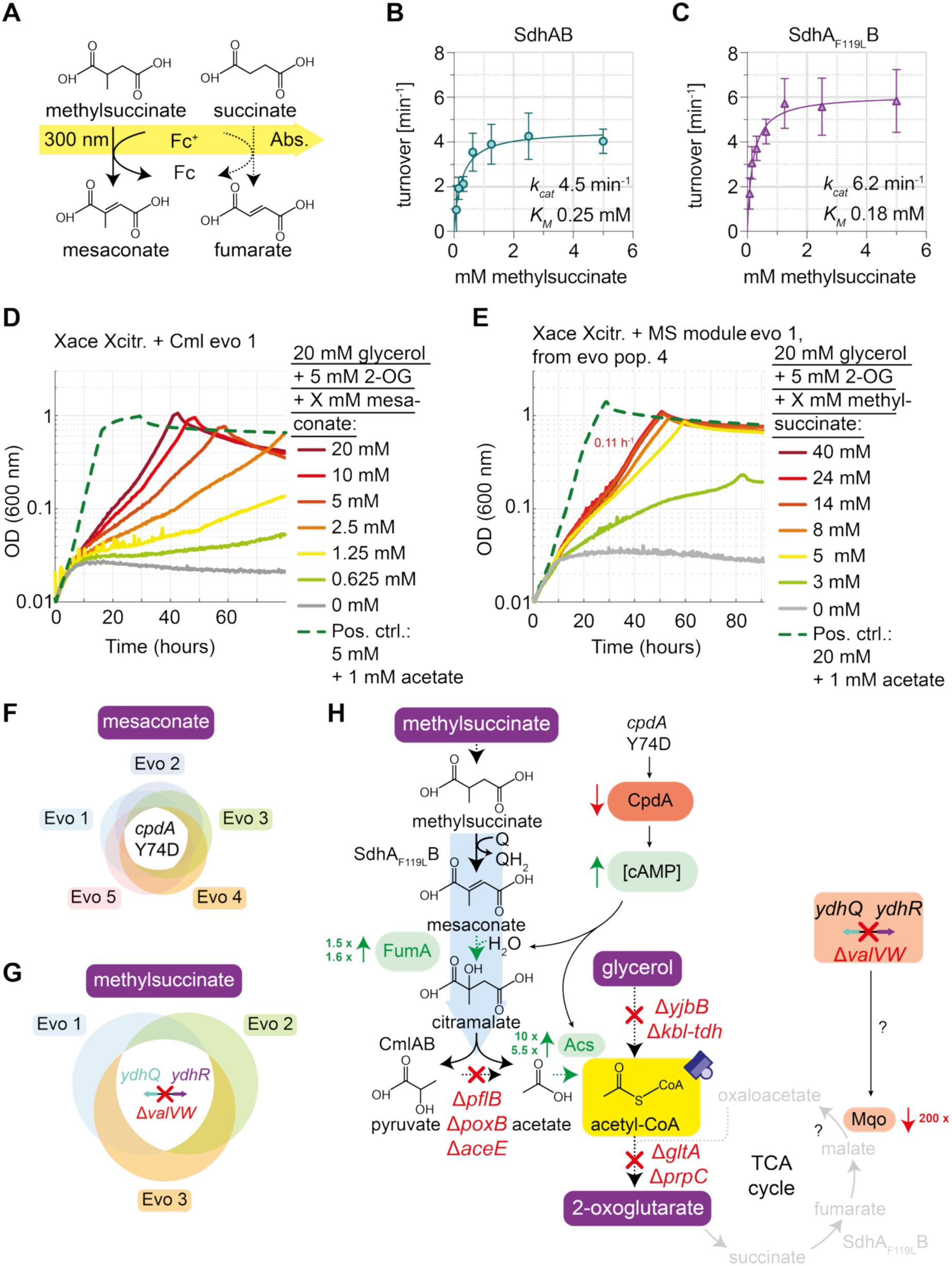
A novel methylsuccinate assimilation module for converts methylsuccinate to pyruvate and acetate. **A)** Assay for *in vitro* succinate dehydrogenase activity measurements. The *E. coli* succinate dehydrogenase natively converts succinate to fumarate (dotted arrow) and promiscuously forms mesaconate from methylsuccinate. Ferrocenium reduction was followed by absorption measurements at 300 nm. **B)** Activity of the wild-type succinate dehydrogenase with methylsuccinate. **C)** Activity of the F119L succinate dehydrogenase variant with methylsuccinate. **D)** A single colony isolate of the evolved Xace Xcitrate strain expressing Cml grows in a mesaconate dependent manner. **E)** Single clone isolate capable of growing with 20 mM glycerol + 5 mM 2-oxoglutarate and varying methylsuccinate concentrations. **F)** All mesaconate evolved isolates share a CpdA Y74D mutation. **G)** All methylsuccinate evolved isolates lack the valine tRNA synthetases *valVW* in addition to the previously acquired CpdA Y74D mutation. **H)** Schematic representation of the hypotheses on how the acquired mutations might affect the endogenous enzymes involved in or surrounding the methylsuccinate assimilation module based on proteomics analyses. After acquisition of the CpdA Y74D mutation, FumA and Acs appear to be upregulated in mesaconate evolved mutants. Following the emergence of the Δ*valVW* mutation in the methylsuccinate evolved strains, *mqo* appears to be strongly downregulated, which could counteract SdhA_F119L_ acting as 2-oxoglutarate sink. A connection between *valVW* and *mqo* expression is not described in literature.

### Host-centric ALE approaches for realizing the methylsuccinate route in vivo

In a next step, we aimed at establishing the methylsuccinate route *in vivo* through adaptation and evolution of the host strain. In these host-centric efforts, several iterative rounds of strain development and testing in combination with experimental evolution and validation are used to realize a synthetic pathway.

To establish the methylsuccinate route, we used a citramalate lyase from *Raoultella planticola* (Cml) that directly cleaves citramalate to pyruvate and acetate (provided as gift from the laboratory of Ivan Berg, Extended Data Figure 3D). We first introduced this enzyme into Pyr-aux. Δ*eno* and tested for growth with RS-citramalate as carbon source but did not observe growth (Extended Data Figure 3E). Speculating that the R-isomer of citramalate might inhibit Cml, we also tested mesaconate as growth substrate, which is converted by endogenous fumarase A exclusively into S-citramalate^27^. However, growth of the Pyr-aux. Δ*eno* was not rescued from mesaconate either (Extended Data Figure 3E), indicating other reasons that interfered with module function. Note that acetate is supplied at 20 mM concentration for energy and biomass formation in the Pyr-aux. Δ*eno* strain. We suspected that these high acetate levels would thermodynamically disfavour the required citramalate cleavage by Cml and interfere with functioning of the methylsuccinate module in the Pyr-aux. Δ*eno* strain.

In the following, we switched to an acetyl-CoA-auxotrophy selection scheme, in parti-cular the Xace Xcitrate strain, which is more than three times as sensitive as the SL2 strain and requires only µM of acetate for growth (Figure 2D, Extended Data Figure 1D). In addition to testing the Cml plasmid tested in the Pyr-aux. Δ*eno* strain, we cloned Cml behind ribosomal binding sites of varying strength and tested for growth with mesaconate or RS-citramalate as acetyl-CoA source (Extended Data Figure 4A). Because the strain did not show initial growth (Extended Data Figure 4B), we decided to perform a semi-relaxing evolution and supplied declining acetate concentrations along with 5 mM mesaconate (Extended Data Figure 4C). During this evolution, one population was raised from the strain expressing the Cml with the weakest rbs ^28^, which was able to grow with mesaconate instead of acetate (Extended Data Figure 4D). Interestingly, RS-citramalate supplementation did not rescue growth of the evolved strain, which indicates either import limitations or inhibition of Cml by R-citramalate (Extended Data Figure 4E). Using single clones isolated from this population, we validated mesaconate- and citramalate-lyase dependent growth for these isolates (Figure 3D, Extended Data Figure 5A; Extended Data Figure 5B, C). Furthermore, we confirmed synthesis of acetyl-CoA from mesaconate by following isotopic label from ^12^C_5_-mesaconate into leucine in presence of U-^13^C-labelled glycerol and 2-oxoglutarate (Extended Data Figure 5D, E).

However, when switching to methylsuccinate as carbon source, the Xace Xcitrate strain expressing citramalate lyase only was not capable of growth (Extended Data Figure 6A), likely due to insufficient methylsuccinate dehydrogenase activity. Thus, we genomically introduced Sdh F119L ^29^ alongside with Cml in the mesaconate evolved background (mesaconate evolved mutant 1, cured of citramalate lyase) and kept the resulting strain (Xace Xcitrate +MS module) in selective medium with methylsuccinate until populations capable of growth emerged (Extended Data Figure 6B). From these populations, we isolated methylsuccinate-dependent single clones, growing robustly at a rate of 0.1 h^-1^, which we further characterized (Figure 3E, Extended Data Figure 6B-E). For these strains, we confirmed incorporation of methylsuccinate-derived carbon into acetyl-CoA (and subsequently leucine) via isotopic labelling (Extended Data Figure 6F, G).

Subjecting the five single clone isolates to whole genome sequencing, we found that all mesaconate-evolved clones carried a Y74D mutation in the cAMP phosphor-diesterase CpdA (Figure 3F, Supplementary Dataset 2), potentially affecting cAMP levels and thus exerting rather broad regulatory effects ^30^. In addition to this mutation, the three methylsuccinate-evolved clones shared a deletion of the GAC-tRNAs *valV* and *valW* (Figure 3G, Supplementary Dataset 3). While *valVW* are not essential to *E. coli*, it is known that their knockout has pleioptropic effects^31^.

To understand the regulatory changes at a systems level, we next analyzed the proteome of the unevolved parent, the mesaconate-evolved strains and the methyl-succinate-evolved descendants (Extended Data Figure 7). When comparing two of the the mesaconate-evolved mutants (evo 1 and 2) with their parent, we found that acetyl-CoA synthetase (Acs), which catalyzes the activation of acetate into acetyl-CoA^32^, was upregulated in all cases (Figure 3H, Supplementary Table 1, Extended Data Figure 7A, B, Supplementary Figure 2A-E). Note that Acs is typically repressed in presence of glycerol (the co-substrate of our selections)^32^ as part of cellular overflow metabolism^33^, and further controlled by cAMP levels^34^. We hypothesized that the CpdA Y74D mutation increased Acs activity in the evolved strains, allowing more efficient conversion of mesaconate-derived acetate into acetyl-CoA, and thus growth (Figure 3H). Indeed, when testing for selective demand, the evolved strains reached higher biomass yields with lower acetate concentrations compared to their parents (Supplementary Figure 2F).

In addition to upregulation of Acs, we also observed upregulation of the cAMP-Crp controlled ^34^ pathway enzyme Fumarase A (Figure 3H, Extended Data Figure 7A, B), likely improving conversion of mesaconate into citramalate^27^. The maltoporin LamB (Extended Data Figure 7A, B) was also upregulated. However, its deletion did not alter growth, indicating rather a non-essential consequence of increased cAMP levels (Extended Data Figure 7C). When comparing the methylsuccinate-evolved isolates to the mesaconate-evolved precursor strains, *mqo* was downregulated in all cases (Extended Data Figure 7D, E and Supplementary Figure 3), indicating a specific adaptation that enabled methylsuccinate-dependent growth. Downregulation of *mqo* probably reduces TCA cycle activity towards oxaloacetate synthesis, thus counteracting the Sdh-dependent drain of 2-oxoglutarate (Figure 3H). In sum, the mutations obtained through experimental evolution appeared to have altered and adopted the native enzyme network of *E. coli* that closely interacts with the synthetic pathway module.

Taken together, focusing on the host strain, our efforts resulted in functional implementation of the methylsuccinate module. Growth on methylsuccinate was achieved in several rounds of experimental evolution of the selection strains and subsequent analysis over the course of six months, resulting in robust growth rates of 0.1 h^-1^. Notably, all adaptations resulted in adjustments of the endogenous metabolic network of the host. However, to what extent these mutations were crucial to establish methyl-succinate-dependent growth, or whether our choice of pathway expression levels made these adjustments essential for growth remained unanswered. To answer this question, we thus attempted to engineer methylsuccinate-dependent growth of an unevolved selection strain through expression level tuning of our methylsuccinate THETA module.

### A module-centric machine-learning guided workflow (MEVIS) to engineer the methyl-succinate route in vivo

Omics characterizations of the evolved strains showed that enzyme expression levels of the native *E. coli* network needed to be fine-tuned to establish the methylsuccinate module, which is a common observation in synthetic pathway engineering^35^. Next, we wanted to take a module-centric approach to assess in how far direct expression level-tuning for Cml and Sdh variant F119L would reduce the need for such host mutations (and thus adaptive laboratory evolution).

To that end we turned our attention to a recent machine learning-guided, semi-automated workflow, METIS, which we had developed mainly for *in vitro* applications, e.g., identification of optimal enzyme and substrate concentrations for *in vitro* networks^36,37^. We adapted this workflow for more efficient solution space sampling and *in vivo* engineering (named MEVIS for **M**achine-learning guided **E**xperimental in **V**ivo **I**mprovement of **S**ystems; Supplementary Note 1, Supplementary Figure 4) and coupled it to a Marionette collection-based^38^ expression system to tune expression levels of individual enzymes in an automated fashion (Figure 4A, B, Supplementary Figure 4, Supplementary Note 1). For our experiments, we tested for optimization of both, enzyme expression levels and carbon source concentrations, and used growth rates multiplied by maximum optical density as readout for growth and thus pathway performance (Figure 4C).

**Figure 4:**
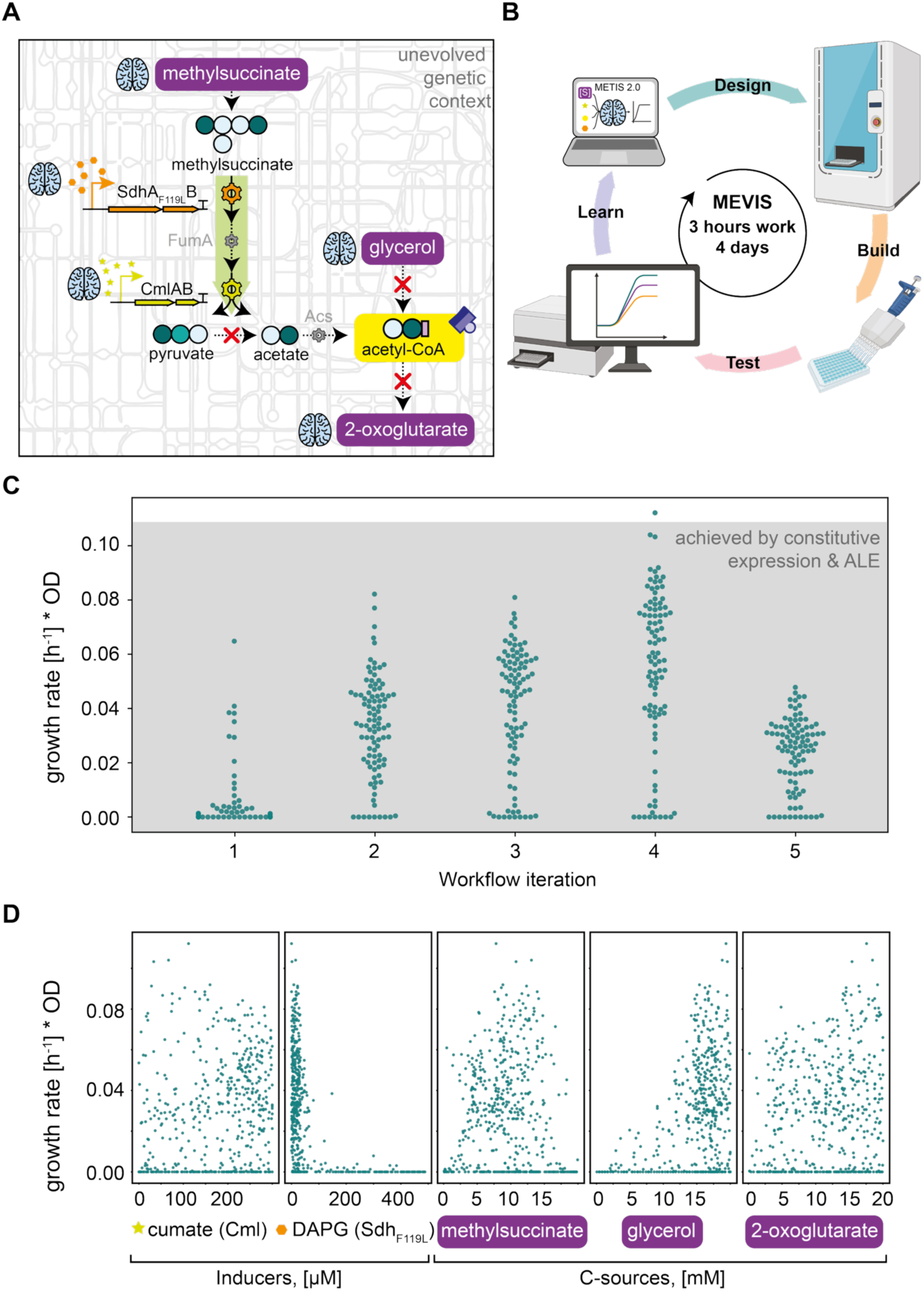
The MEVIS workflow allows engineering methylsuccinate assimilation without adaptive laboratory evolution. **A)** Schematic representation of the methylsuccinate assimilation module improved by MEVIS. Inducer concentrations for citramalate lyase and succinate dehydrogenase (Sdh_F119L__B) expression and carbon source concentrations (glycerol, 2-oxoglutarate, methylsuccinate) were optimized in five workflow rounds. **B)** The MEVIS workflow combines outputs of the optimized METIS model with automated pipetting using an ECHO liquid handling robot. Cells are added to multi-well plates after medium preparation, followed by growth monitoring for four days. **C)** Four rounds of MEVIS based optimisation helped find the growth conditions allowing maximal growth (indicated by growth rate multiplied with OD). The grey box indicates growth reached with adaptive laboratory evolution only. Each round was performed in n=3 replicates, of which the average is shown. **D)** Dependency of growth (growth rate multiplied with OD) on inducer and carbon source concentrations (averages of n=3 experiments are shown). Succinate dehydrogenase expression levels converge to low µM concentrations.

Notably, already the first round of MEVIS identified multiple combinations of enzyme expression levels and carbon source concentrations that allowed the unevolved strain to directly grow with methylsuccinate (Figure 4C, Extended Data Figure 8), indicating that expression tuning of the module can – to some extent – overcome the need for adaptive laboratory evolution of the host. Interestingly, these tuned strains showed growth rates (0.1 h^-1^) that were comparable to that of the methylsuccinate-evolved strain (Extended Data Figure 8A, Figure 3E), while their final optical densities were (still) lower than those of the evolved strain (Extended Data Figure 8B). However, with each MEVIS iteration, overall growth was optimized over four workflow rounds across a time span of three weeks (Figure 4C, Extended Data Figure 8). Overall, these findings showed that module-centric optimization can result in similar outcomes as experimental evolution of the host, albeit in much shorter time.

When globally analyzing the data from all MEVIS rounds to identify common trends, we observed that growth appeared to be constrained to very low to basal induction levels of the Sdh F119L (Figure 4D, Extended Data Figure 8C). At the same time, both yields and cell densities benefited from supplying maximum quantities (20 mM) of glycerol and 2-oxoglutarate, whereas the methylsuccinate concentration appeared to be optimal at around 10 mM (Figure 4D, Extended Data Figure 8C). Since growth rates did not increase further after the first round of MEVIS and the optimal methylsuccinate concentration was lower than the one tolerated by the evolved strain (Figure 3E), we speculated that flux through the methylsuccinate assimilation module was potentially constrained by FumA and Acs availability. Since we suspected that the downregulation of *mqo* in the methylsuccinate evolved strains might not improve pathway flux but would merely compensate overly strong expression of succinate dehydrogenase, we sought to apply MEVIS based module optimization in the mesaconate-evolved host, in which Acs and FumA expression had been increased through experimental evolution.

### Combining the evolved genetic context with module optimisation through MEVIS allows optimal strain growth with methylsuccinate

Having shown that host-centric and module-centric approaches both resulted in similar phenotypic outcomes, albeit through separate mechanisms, we became interested to what extent combination of both approaches could further improve methylsuccinate-dependent growth (Figure 5A). Thus, we used the mesaconate-based evolved strain as background and optimized the methylsuccinate module through MEVIS. Round 1 of MEVIS directly resulted in several tuned strains that grew on methylsuccinate. Further rounds of MEVIS improved both growth rate and biomass formation significantly compared to the unevolved strain (Figure 5B, C). Since at high optical densities we wanted to prioritize enhancing pathway turnover over yield, we constrained optical densities to a maximum upper input value of 1.5 during the optimization (Extended Data Figure 9A). The optimized evolved strain grew at higher induction levels for both Cml and the mutant Sdh as before and tolerated higher methylsuccinate concentrations (Figure 5D), which indicated that the evolved genetic background allowed for more flux through the methylsuccinate assimilation module. This was further supported by the fact that growth rates continuously improved with each MEVIS round – in contrast to MEVIS-based optimization efforts in the background of the unevolved strain, which had stalled at 0.1 h^-1^ (Extended Data Figure 9A). In addition, the biomass yield also improved significantly from MEVIS round 1 to MEVIS round 4 (Extended Data Figure 9A), which resulted in growth parameters that were 200% improved (with a maximum growth rate of 0.19 h^-1^ and maximum OD of 1.5) compared to either the evolved strain (maximum growth rate of 0.1 h^-1^ and maximum OD of 1.1) or the MEVIS-tuned strain alone (maximum growth rate of 0.1 h^-1^ and maximum OD of 1.2) (Figure 5E, Extended Data Figure 9A, B). Notably, while MEVIS-optimization resulted in immediate and improved growth of the mesaconate-evolved background, achieving growth with fixed expression levels of Cml and Sdh F119L still required further (short-term) evolution (Extended Data Figure 6).

**Figure 5:**
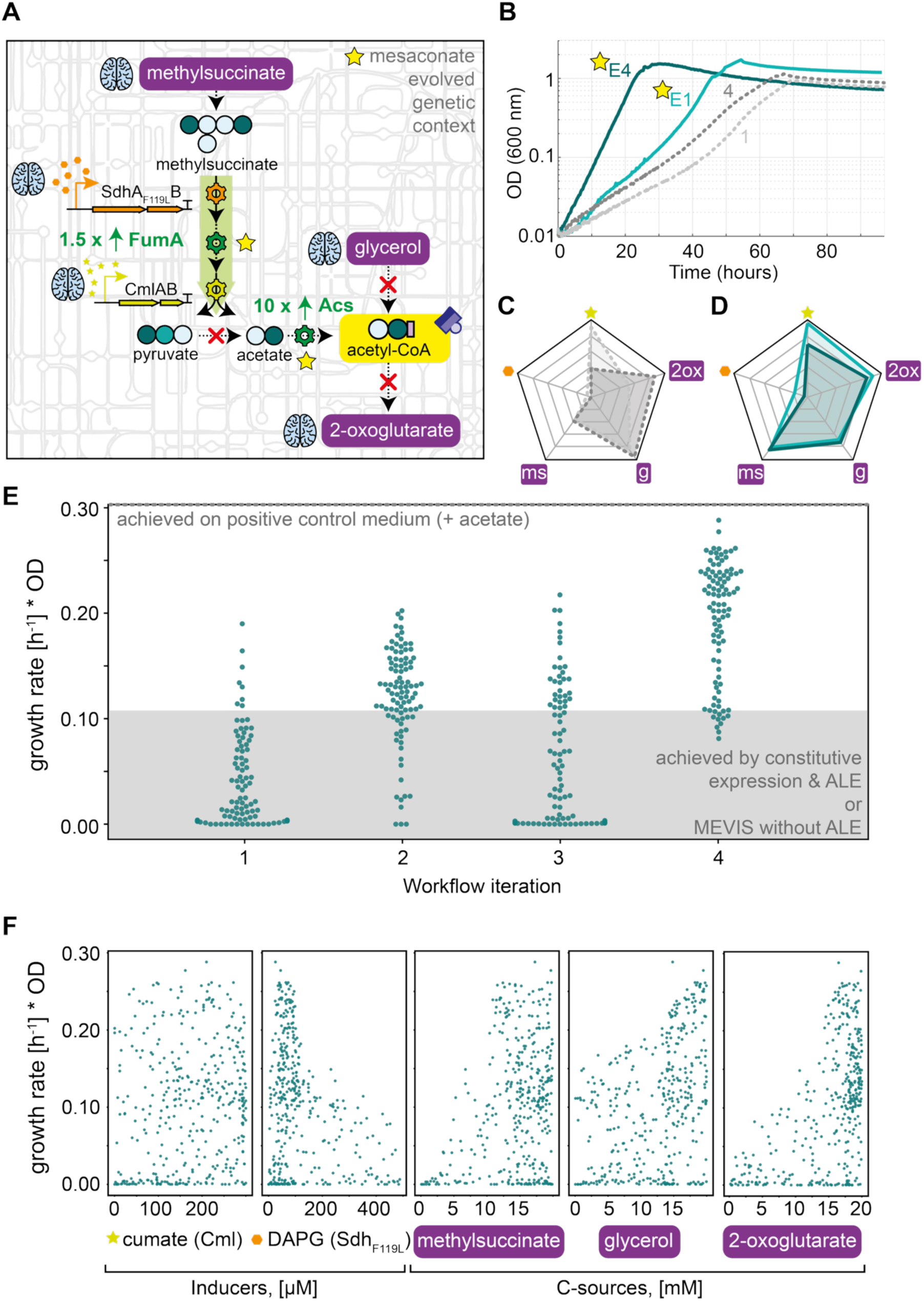
Combining MEVIS and an evolved genetic background yields an optimized methylsuccinate dependent strain. **A)** Schematic representation of the methylsuccinate assimilation module improved by MEVIS. Inducer concentrations for citramalate lyase and succinate dehydrogenase (Sdh_F119L__B) expression and carbon source concentrations (glycerol, 2-oxoglutarate, methylsuccinate) were optimized in four workflow rounds. The mesaconate evolved strain background with upregulated FumA and Acs was used as chassis to optimise methylsuccinate dependency. **B)** Cell growth of the best test condition of MEVIS rounds one and four in the unevolved parental strain (grey curves) and with the mesaconate evolved genetic background (teal and turquoise curve, evolved background indicated by stars). While improvements between the best condition of round one and round 4 are minute in the unevolved background, the evolved context permitted growth rate and biomass yield improvements between the best conditions of round one and round 4. **C)** Radar chart displaying the differences between the optimal concentration combinations of round one (light grey) and round four (dark grey) of the unevolved strain. Concentrations at the chart center are zero, the corner point concentrations are defined as follows: cumate (lime star) = 300 µM; DAPG (orange hexagon) = 500 µM; methylsuccinate (ms), glycerol (g), 2-oxoglutarate (2ox) = 20 mM. **D)** Radar chart displaying the differences between the optimal concentration combinations of round one (turquoise) and round four (teal) of the mesaconate evolved strain background. Concentrations at the chart center are zero, the corner point concentrations are defined as follows: cumate (lime star) = 300 µM; DAPG (orange hexagon) = 500 µM; methylsuccinate (ms), glycerol (g), 2-oxoglutarate (2ox) = 20 mM. **E)** Four rounds of MEVIS based optimisation helped find the growth conditions allowing optimal strain growth (indicated by growth rate multiplied with OD). The grey box indicates growth reached with adaptive laboratory evolution only. The dotted grey line indicates growth reached in positive control medium (with 1 mM acetate). Each round was performed in n=3 replicates, of which the average is shown. **F)** Dependency of growth (growth rate multiplied with OD) on inducer and carbon source concentrations (averages of n=3 experiments are shown).

Data analysis showed that mesaconate-dependent growth was promoted by low expression levels of Sdh F119L (Figure 5F, Extended Data Figure 9C, D). Yet, these levels were significantly higher compared to those of the MEVIS-tuned module in the unevolved background, indicating higher flux capacity of the evolved strain (Figure 4D). Along these lines, optimal growth of the evolved strain was shifted to higher methylsuccinate concentrations (Figure 5F, Extended Data Figure 9C, D) supporting increased flux through the module and fast higher growth. Ultimately, these findings showed that combination of experimental evolution and MEVIS-based module optimization was capable to overcome the limitations of methylsuccinate-dependent growth.

## Discussion

Although synthetic metabolism offers new possibilities for radically new biochemical transformations, there is a major gap between the theoretical design of new-to-nature pathways and their *in vivo* realization. This is apparent for synthetic CO_2_-fixation pathways and in particular for Ccr-based designs, which have been demonstrated *in vitro*^3,4,9,20^, but whose implementation in microbial hosts remains challenging^8^. In this work, we developed suites of pyruvate and acetyl-CoA auxotrophic *E. coli* sensor strains that could be employed to adopt pathway modules revolving around Ccr into the native metabolic network of *E. coli*. As a proof-of-principle, we show successful implementation of the methylsuccinate module of the THETA cycle, advancing our capabilities towards establishing complete orthogonal synthetic CO_2_-fixation pathways in the future.

Notably, our study used two different approaches to achieve implementation of the methylsuccinate module. We show that host-centric, as well as module-centric approaches both lead to basic growth on methylsuccinate. While host-centric approaches required several DBTL rounds and experimental evolution over several months, our pathway-centric approaches guided by machine learning (MEVIS) achieved similar output within three weeks. This indicates that pathway module tuning can to some extent bypass – or more precisely complement – the need for experimental evolution to adjust fluxes between native metabolism and a synthetic module, enabling (initial) strain growth. Although we note that for highly efficient implementation, experimental evolution can still be an important and necessary factor.

Our findings fall in line with other implementation efforts in *E. coli*, such as the Calvin cycle^15,23^, the reductive glycine pathway^10^, the CORE cycle^39^, the serine threonine cycle^11^, or the EuMP cycle^16^. These examples all showed that successful *in vivo* transfer required synchronization of the endogenous metabolic network with the synthetic module, which was achieved via adaptive laboratory evolution. However, their implementation was achieved by exclusively focusing on strain adaptation. Through MEVIS, such adjustments can be performed now more systematically, from the pathway-side and in parallel or in combination with evolution of the host strain.

The combination of machine learning-guided experimentation with automated pipetting has been successfully applied to optimize different biological systems, including cell-free assays^9,36^ and microbial cell factories^40^ ^41^. Methodologically, these efforts could be further improved by extending the platform expression system to accommodate larger arrays of inducible operons and combining tuning of pathway enzyme expression with inducible knockdowns of surrounding endogenous reactions, for example using CRISPRi-based approaches^41–44^. Thus, combining growth-coupled selections with machine-learning guided experimental design and lab automation holds great potential to accelerate and improve the design-build-test-learn cycles for metabolic engineering and synthetic metabolism in the future.

## Methods

### Protein production and purification

To produce recombinant succinate dehydrogenase variants, *E. coli* BL21 (DE3) (for all proteins except the mutases) were transformed via heat-shock with the pCDFDuet-SdhAB expression plasmids and plated on LB Spec plates. The cells were scraped off the plate and inoculated in TB (12 g/L tryptone, 24 g/L yeast extract, 17 mM KH2PO4, 72 mM K2HPO4, 0.5% (v/v) glycerol) with antibiotic. The cells were grown at 37°C, 120 rpm until an OD600nm of ∼0.7-0.9 was reached. The cultures were cooled down to 25°C (BL21(DE3)) and induced with 500 μM IPTG. The culture was incubated overnight and harvested the next day (4,000 xg, 10 min, 10°C). The pellet was resuspended in Buffer A (50 mM HEPES pH 7.5, 500 mM KCl) and sonicated on ice to lyse the cells (KE76 probe, 1 min, 01s on 01s off, 3x in total). The lysed cells were centrifuged (100,000 g, 45 min, 4°C) to separate soluble and insoluble fraction. The soluble fraction was loaded onto Ni-NTA agar resin (Macherey-Nagel) that was previously equilibrated with wash buffer (50 mM HEPES pH 7.5, 75 mM Imidazole, 500 mM KCl). The resin was washed with 70 mL wash buffer and then eluted with 2 mL Buffer B (50 mM HEPES pH 7.5, 500 mM Imidazol, 500 mM KCl). The proteins were desalted into Buffer D (50 mM HEPES/KOH pH 7.5, 150 mM KCl) using PD-10 columns (Cytiva) and concentrated on Amicon filters. The proteins were flash frozen in liquid N2 and stored at −70°C.

### *In vitro* characterization of Succinate dehydrogenase

To determine the activity of succinate dehydrogenase, the absorbance change due to ferrocenium reduction was monitored at 300 nm (Supplementary Figure 2A). For this, 0.25 mM ferrocenium hexafluorophosphate, 100 mM MOPS pH 7.5, 1 µM enzyme and varying concentrations of succinate or methylsuccinate were mixed in a 200 µl High Precision Cell 10 mm Light Path quartz cuvette (Hellma Analytics, Müllheim, Germany). The reduction of ferrocenium upon substrate addition was followed by measuring the change of absorption at 300 nm in a Cary60 UV-Vis spectrophotometer (Agilent, Santa Clara, CA, USA) at 30 °C with the kinetics program with 0.5 s save time. Initial velocity measurements were performed in triplicates, and turnover frequencies were calculated based on Lambert-Beer’s law with ε(300 nm) = 4.3 mM^-1^ cm^-1^ and fit using the Michaelis-Menten function in Prism 8 (Graphpad, San Diego, California, USA).

### Strain construction

All *E. coli* strains used in this study are listed in Supplementary Table 2. We used strain SIJ488 as a wild type, which carries inducible recombinase and flippase genes^45^. Gene deletions were done by either λ-red recombineering or P1-transduction as described below.

### Gene deletion and genome integration by recombineering

To delete genes by λ-red recombineering, chloramphenicol resistance cassettes with overhangs homologous to the target locus were generated by PCR using KO primers (Supplementary Table 3), the chloramphenicol (Cap) cassette from pKD3 (pKD3 was a gift from Barry L. Wanner (Addgene plasmid #45604; http://n2t.net/addgene:45604; RRID:Addgene_45604)) as template and PrimeStar Max polymerase (Takara Bio, Saint-Germain-en-Laye, France). For gene deletion, the target strains were inoculated in LB and grown to OD ∼0.4-0.5, followed by addition of 15 mM L-arabinose to induce recombinase gene expression during 45 min cultivation at 37°C. Then, cells were harvested and washed three times with ice cold 10% glycerol (11,000 rpm, 30 sec, 4°C). Electroporation was performed using ∼300 ng of Cap resistance cassette and 100 µL washed cells (1 mm cuvette, 1.8 kV, 25 µF, 200 Ω). To select for successful gene deletion, cells were plated on LB with the relevant antibiotics. Further deletion confirmation was done by PCR, and antibiotic resistance removal was performed by growing cells to OD ∼0.3, inducing flippase expression by addition of 50 mM L-rhamnose, cultivation overnight at 30 °C and subsequent isolation of single colonies on LB plates. Successful marker removal was confirmed by testing for antibiotic sensitivity and by PCR on the respective locus. To chromosomally integrate citramalate lyase and succinate dehydrogenase (*sdhA_*F119L *sdhB*), the genes were initially cloned into pKD3 (see section “Plasmid construction” below) and subsequently amplified with PrimeStar Max polymerase and primers carrying 50 bp overhangs to the SS9 locus. To remove residual PCR template, the PCR product was digested with DpnI (FastDigest, Thermo Scientific) before purification. The purified PCR product was introduced into the genome via λ-red recombineering and successful integration was selected for via antibiotics as described above for deletions.

### Gene deletion via P1 transduction

Deletions of *gcl*, *aceE*, *pgk, mgsA, tesB, eno* and *gapA* to construct the pyr-aux strain and were performed by P1 phage transduction (Thomason et al, 2007). For *gcl*, *aceE*, *mgsA* and *tesB* deletion, the donor strains carrying single gene deletions with a kanamycin-resistance gene (KmR) as selective marker were taken from the Keio collection (Baba et al, 2006). For *pgk, eno* and *gapA* deletion, the previously constructed single deletion SIJ488 strains with chloramphenicol (CapR) resistance were used as donor strains. To select for successful gene deletion after transduction, the strains were plated on medium with appropriate antibiotics and 20 mM of sodium citrate for P1 phage removal. Single colonies were restreaked once on another plate with antibiotic and citrate. For these, gene deletion was further confirmed by determining the size of the respective genomic locus *via* PCR using KO-Ver primers (Supplementary Table 3) and DreamTaq polymerase (Thermo Scientific, Dreieich, Germany). To remove the antibiotic resistance gene, a fresh culture of the deletion strain was grown to OD_600_ ∼ 0.2, where 50 mM L-rhamnose were added to remove flippase expression during the following cultivation at 30°C for ∼4h. Antibiotic resistance removal was confirmed by testing antibiotic sensitivity of individual colonies on LB with antibiotic compared to LB only, and by PCR for the respective locus using KO-Ver primers and DreamTaq polymerase (Thermo Scientific, Dreieich, Germany).

### Plasmid construction

Cloning was planned using Snapgene (GSL Biotech LLC, San Diego, US). For gene overexpression, Cml (citramalate lyase from *Raoultella planticola* was cloned into a pZ-ASS vector with strong promoter and ribosome binding site C for each subunit. The genes were a gift from Ivan Berg. Cloning was performed in *E. coli* DH5α. For genome integration, the genes were transferred from pZ-ASS to pKD3 by using the NEB HiFi assembly kit (New England Biolabs), which involved amplifying the inserts individually using PrimeStar Max, followed by DpnI digest and purification of fragments for subsequent HiFi assembly. HiFi assemblies were performed using *E. coli* NEB5α cells. Correct insert sizes were confirmed using DreamTaq polymerase (Thermo Scientific, Dreieich, Germany) and primers pZ-ASS-F and pZ-ASS-R for pZ-ASS and pKD3-F and pKD3-R for pKD3. To inducibly express the THETA cycle genes, we first introduced them to the Marionette collection plasmids pAJM.712 (citramalate lyase, for expression in strains carrying the genome repressor array = pSM87), pAJM.657 (citramalate lyase, repressor on plasmid = pSM99), pAJM.711 (succinate dehydrogenase, for expression in strains carrying the genome repressor array = pSM94), and pAJM.847 (succinate dehydrogenase, repressor on plasmid = pSM100) using the HiFi assembly kit provided by New England Biolabs, Ipswich, MA and primers HSM26_pAJM.712_marion_rvs and HSM25_pAJM.712-CML_marion_fwd to amplify Marionette collection plasmids, HSM27_CML_marion_fwd and HSM28_CML_marion_rvs to amplify citramalate lyase and HSM-45_SdhA_Mar_fwd and HSM-46_SdhB_Mar_rvs to amplify succinate dehydrogenase. To co-express multiple genes inducibly from plasmids with compatible origins of replication and different antibiotic resistances, inducible expression vectors from the Marionette collection ^38^ were made compatible with the SEVA collection ^46^ by introducing PacI and SpeI sites relevant to clone the cargo unit (promoter and gene of interest) into SEVA vectors. For this, we introduced PacI and SpeI sites into pSM100 using primers HSM26_pAJM.712_marion_rvs and HSM25_pAJM.712-CML_marion_fwd, and introduced the inducible Sdh operon into a pSEVA624 backbone by restriction of donor and recipient vectors with PacI SpeI, gel purification of the relevant fragments and subsequent ligations (yielding pSM148). Successful assembly of pSM148 was confirmed by colony PCR. The sequence of vectors with correct insert size was confirmed by Sanger sequencing (LGC Genomics, Berlin, DE or Microsynth Seqlab, Göttingen, DE). For in silico sequence analysis, the software Snapgene was used. Target strains were transformed with 40 ng of the correct plasmids by electroporation in the same manner described for transformations with λ-red knockout cassettes. Successful transformation was confirmed by resistance to plasmid specific antibiotics as well as colony PCR with plasmid specific primers and DreamTaq polymerase (Thermo Scientific, Dreieich, Germany).

### Media and growth experiments

LB medium (1% NaCl, 0.5% yeast extract, 1% tryptone) was used for strain maintenance, cloning and deletion strain construction. Pyr-aux. strains with glycolysis disruptions were cultivated in MX medium (M9 medium with 5 g/L casein hydrolysate and 20 mM glycerol + 20 mM succinate) instead of LB medium at all times. For any acetyl-CoA auxotrophic strains, 1 mM acetate was supplied to the medium. If required, antibiotics (kanamycin (50 μg/mL), ampicillin (100 μg/mL), streptomycin, (100 μg/mL), chloramphenicol (30 μg/mL)), or spectinomycin (50 μg/mL) were added. Antibiotics were omitted in growth experiments. Standard M9 minimal media (50 mM Na2HPO4, 20 mM KH2PO4, 1 mM NaCl, 20 mM NH4Cl, 2 mM MgSO4 and 100 μM CaCl2, 134 μM EDTA, 13 μM FeCl3·6H2O, 6.2 μM ZnCl2, 0.76 μM CuCl2·2H2O, 0.42 μM CoCl2·2H2O, 1.62 μM H3BO3, 0.081 μM MnCl2·4H2O) was used for growth experiments and strain evolution with the carbon sources indicated in the text. For growth experiments, precultures were prepared in M9 medium supplemented with and antibiotics for any plasmids present and carbon sources depending on the strain. For strains expressing crotonate utilization genes appropriate inducers and 1 µM B12 (cyanocobalamine) were added to all growth media. After harvesting grown overnight cultures (6,000 xg, 3 min), the cells were washed three times in M9 medium to remove residual carbon sources, antibiotics and cofactors. Growth experiments were performed in 96-well microtiter plates (Nunclon Delta Surface, Thermo Scientific) at 37°C and were inoculated with washed cells to an optical density (OD600) of 0.01 in 150 µL total culture volume per well. To avoid evaporation but allow gas exchange, 50 μL mineral oil (Sigma-Aldrich) were added to each well. If not stated otherwise, growth was monitored in technical triplicates in a BioTek Epoch 2 Microtiterplate reader (BioTek, Bad Friedrichshall, Germany) by absorbance measurements (OD600) of each well every ∼10 minutes with intermittent orbital and linear shaking. As previously established empirically for the instrument, blank measurements were subtracted and OD600 measurements were converted to cuvette OD600 values by multiplying with a factor of 4.35. Growth curves represent the average of technical triplicate measurements and were plotted in MATLAB version R2020a.

### 13C-labelling of proteinogenic amino acids

For labelling analysis of the evolved Xace Xcitrate strains growing with mesaconate or methylsuccinate, the evolved strains were cultured in 3 ml M9 minimal medium supplemented with 20 mM [U-^13^C] glycerol, 5 mM [U-^13^C] glutamate and 20 mM mesaconate or methylsuccinate. At the late exponential phase, cells with the amount equivalent to 1 ml OD600 of 1 were collected, washed with water once and resuspended in 1 ml 6 M HCl to hydrolyse the biomass at 95 °C overnight. Then, the samples were completely dried under an airstream at 95 °C, re-dissolved in 1 ml H2O and centrifuged to remove insoluble particles. Amino acids (^13^C-labeled amino acid) were analyzed using HRES LC-MS, performing the chromatographic separation on an Agilent Infinity II 1260 HPLC system with a ZicHILIC SeQuant column (150 × 2.1 mm, 5 µm particle size, 100 Å pore size) connected to a ZicHILIC guard column (20 × 2.1 mm, 5 µm particle size) (Merck KgAA) at a constant flow rate of 0.3 ml/min. Mobile phase A was 0.1 % formic acid in 99:1 water:acetonitrile (Honeywell) and phase B was 0.1 % formic acid in 99:1 acetonitrile:water. The column oven was set to 25 °C, the autosampler was maintained at 4 °C. An injection volume of 1 µl was used. The mobile phase profile consisted of the following steps and linear gradients: 0 to 12 min from 90 to 70% B; 8 12 to 15 min from 70 to 20% B; 15 to 17 min constant at 20% B; 17 to 17.1 min from 20 to 90% B; 17.1 to 20 min constant at 90% B. An Agilent 6550 ion funnel Q-TOF mass spectrometer was used in positive mode with a duel jet stream electrospray ionization source and the following conditions: ESI spray voltage 1000 V, nozzle voltage 900 V, sheath gas 200° C at 9 L/min, nebulizer pressure 35 psig, and drying gas 130° C at 12 L/min. The TOF was calibrated using an ESI-L Low Concentration Tuning Mix (Agilent) before measurement (residuals less than 2 ppm for five reference ions) and was recalibrated during a run using 121.050873 m/z as reference mass. MS data were acquired with a scan range of 50-250 m/z. LC-MS data were analyzed using MassHunter Qualitative Analysis software (Agilent).

### MDF analysis

To determine the net reaction equations of all acetyl-CoA assimilation modules and evaluate their thermodynamic feasibility in physiological conditions, Max-Min Driving Force (MDF) analysis was conducted using the Python packages equilibrator API and equilibrator pathway (version 0.6.0 for both) ^47^. Changes in Gibbs Free Energy were estimated using the Component Contribution Method ^47^. For all carboxylation reactions, CO_2_ (rather than bicarbonate) was considered as substrate since its concentration is pH-independent ^9^. Default values were used, i.e. the pH was set to 7.5, magnesium concentration was set to 1 mM (pMg = 3), ionic strength 0.25 M, and metabolite and cofactor concentrations were constrained to physiologically relevant concentrations (1 µM – 10 mM) as described previously ^48^.

### Sequence analysis of evolved strains

For whole genome sequencing, strains were grown overnight in LB medium supplemented with appropriate antibiotics. The Macherey-Nagel NucleoSpin Microbial DNA purification Kit (Macherey-Nagel, Düren, Germany) was used to extract the genomic DNA. Microbial short insert PCR-free library construction for single-nucleotide variant detection and generation of 150 bp paired-end reads on an Illumina HiSeq 3000 platform were performed by Novogene (Cambridge, UK). Breseq (Barrick Lab, Texas)^49^ was used to map the obtained reads to the reference genome of *E.coli* MG1655 (GenBank accession no. U00096.3). With the algorithms supplied by the software package, we identified single-nucleotide variants (with >50% prevalence in all mapped reads) and regions more than 2 standard coverage deviations from the global median coverage.

### MEVIS workflow for the METIS-guided optimisation of expression levels *in vivo*

To optimize enzyme expression levels and carbon sources, inducer and carbon source combinations were pipetted using ECHO 650 and ECHO 525 liquid handling robots. Combinations to test were defined by the METIS algorithm that was updated to use Hammersley sampling instead of random sampling in the first workflow round. Before preparing the experiment, the ECHO liquid handlers were calibrated according to the manufacturer protocols. Stock solutions of cumate (100 mM), diacetylphloroglucinol (DAPG, 25 mM) and glycerol (2 M) were pipetted in 58 µl volumina into a Beckman Coulter 384 well plate (source plate) and transferred according to the METIS instructions to a 96 well Thermo scientific flat bottom Nunclon Delta surface recipient plate by an ECHO 625. Next, stock solutions of 2-oxoglutarate (500 mM), methylsuccinate (200 mM) and M9 minimal medium were pipetted in a Beckman Coulter 6 well reservoir plate to be subsequently transferred into the 96 well plate already containing the inducer combinations and glycerol using an ECHO 525. To be able to inoculate the experiment at a defined optical density, each well was filled to a total volume of 35 µl with M9 medium by the ECHO 525 robot. Test cultures were washed as described for growth experiments and then used to inoculate the experiment at a starting OD of 0.01 and a total volume of 150 µl (i.e., 115 µl of culture normalized to OD 0.013 were added to the 35 µl in the 96 well plate). Mineral oil was added as described for growth experiment. For each round, three replicate 96 well plates were prepared and monitored in three separate BioTek plate readers. Growth experiments were analysed using MATLAB as described above, and the growth readouts of final OD and growth rate were compiled and averaged in Excel, to feed the average values for each combination back to METIS for the generation of the next test combinations. To ensure the reliability of the cultivation, we confirmed the correct strain genotype to be present in the five best-growing wells of round four in each MEVIS application by confirming all deletions and presence of all plasmids by colony PCR.

## Author contributions

H. S.-M. and T. J. E. conceptualized the study. H. S.-M., K. S., T. D., H. K., A. S., H. H., E. R., S. H. L., M. K., and B. D. constructed the strains of this study. K. S. optimized the METIS algorithm for in vivo applications. H. S.-M. designed and realized the MEVIS workflow with assistance from N. B. J. K. and T. G. analysed proteomics samples, P. C. and N. P. analysed LC-MS data. V. R. developed the Sdh mutant variant, which H. S.-M. tested in vitro. S. L. created plasmids used in this study and designed the methylsuccinate assimilation module. H. S.-M. and T. J. E. wrote the manuscript with input from V. R., S. H. L., B. D., and N. J. C. N. J. C. and T. J. E. supervised the research.

## Acknowledgements

This work was supported by the Max Planck Society and the the Bosch Research Foundation. H. S.-M. acknowledges funding by the Bosch Research Foundation and the Joachim Herz Foundation (in form of an Add-on Fellowship for interdisciplinary life sciences).

## Extended Data Figures

**Extended Data Figure 1:**
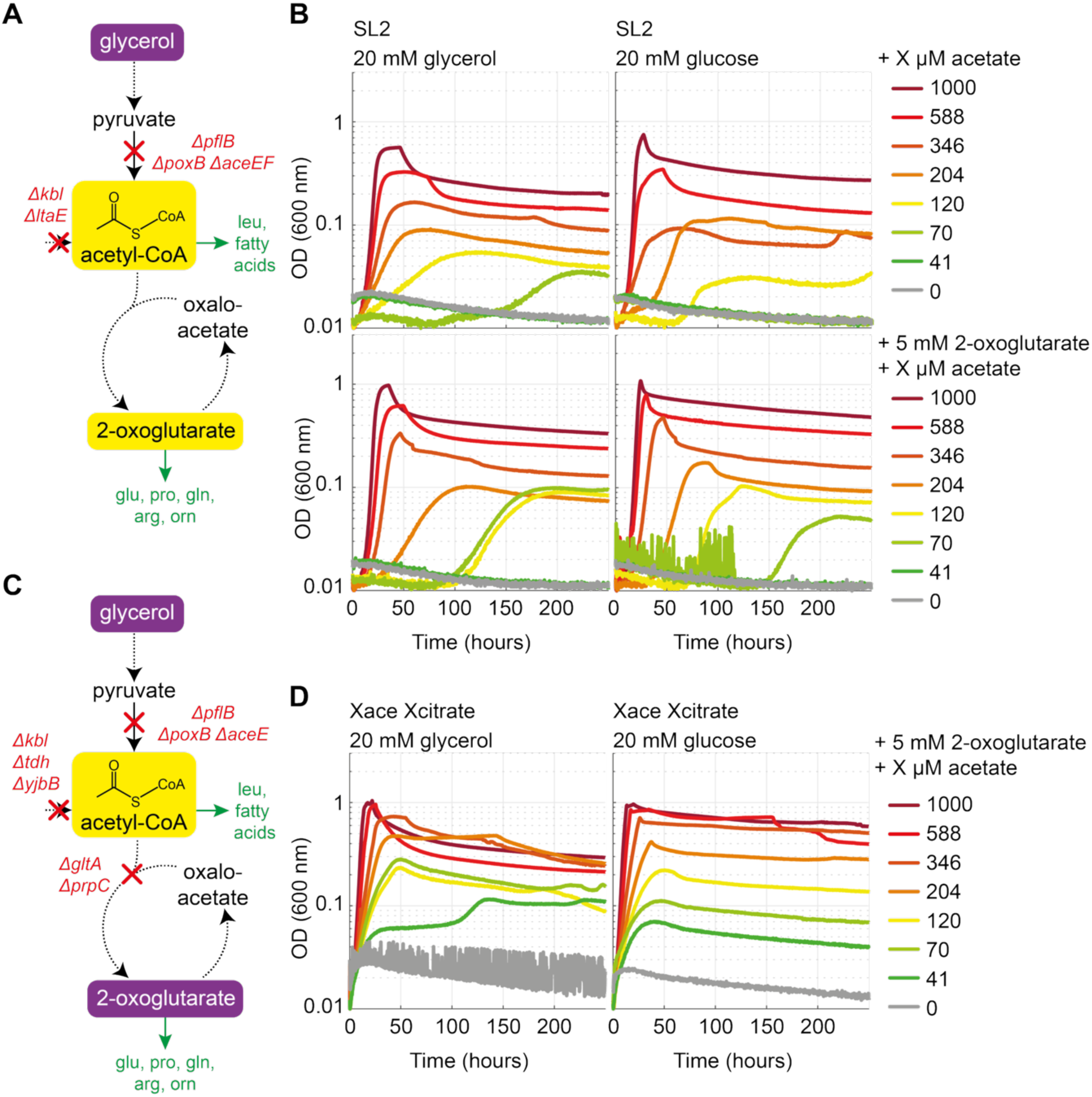
Construction and characterisation of acetyl-CoA auxotrophic sensor strains. **A)** Selection scheme for the SL2 strain created by Luo et al^9^. Acetyl-CoA is metabolically isolated from pyruvate by deletions of pyruvate dehydrogenase (*aceEF*), pyruvate-formate lyase (*pflB*), pyruvate oxidase (*poxB*). The secondary acetyl-CoA sources 2-amino-3-ketobutyrate CoA ligase (*kbl*) and threonine dehydrogenase (*tdh*) are deleted. Acetyl-CoA is required to synthesize leucine, fatty acids and TCA cycle intermediates. **B)** Growth of the SL2 strain in presence of varying acetate concentrations on glycerol (left) or glucose (right) without (top) and with (bottom) 2-oxoglutarate supplementation. **C)** Selection scheme for the Xace Xcitrate strain. The selection strain was constructed in an SIJ488 background by deleting the acetyl-CoA sources deleted in the SL2 strain (Δ*pflB* Δ*poxB* Δ*aceE* Δ*kbl* Δ*tdh*) and removing the citrate synthase (Δ*gltA* Δ*prpC*) as major metabolic acetyl-CoA sink. Due to the deletion of citrate synthase, the strain relies on external 2-oxoglutarate supplementation. **D)** Growth of the Xace Xcitrate strain with glycerol or glucose as main carbon source in presence of 5 mM 2-oxoglutarate and decreasing acetate concentrations.

**Extended Data Figure 2:**
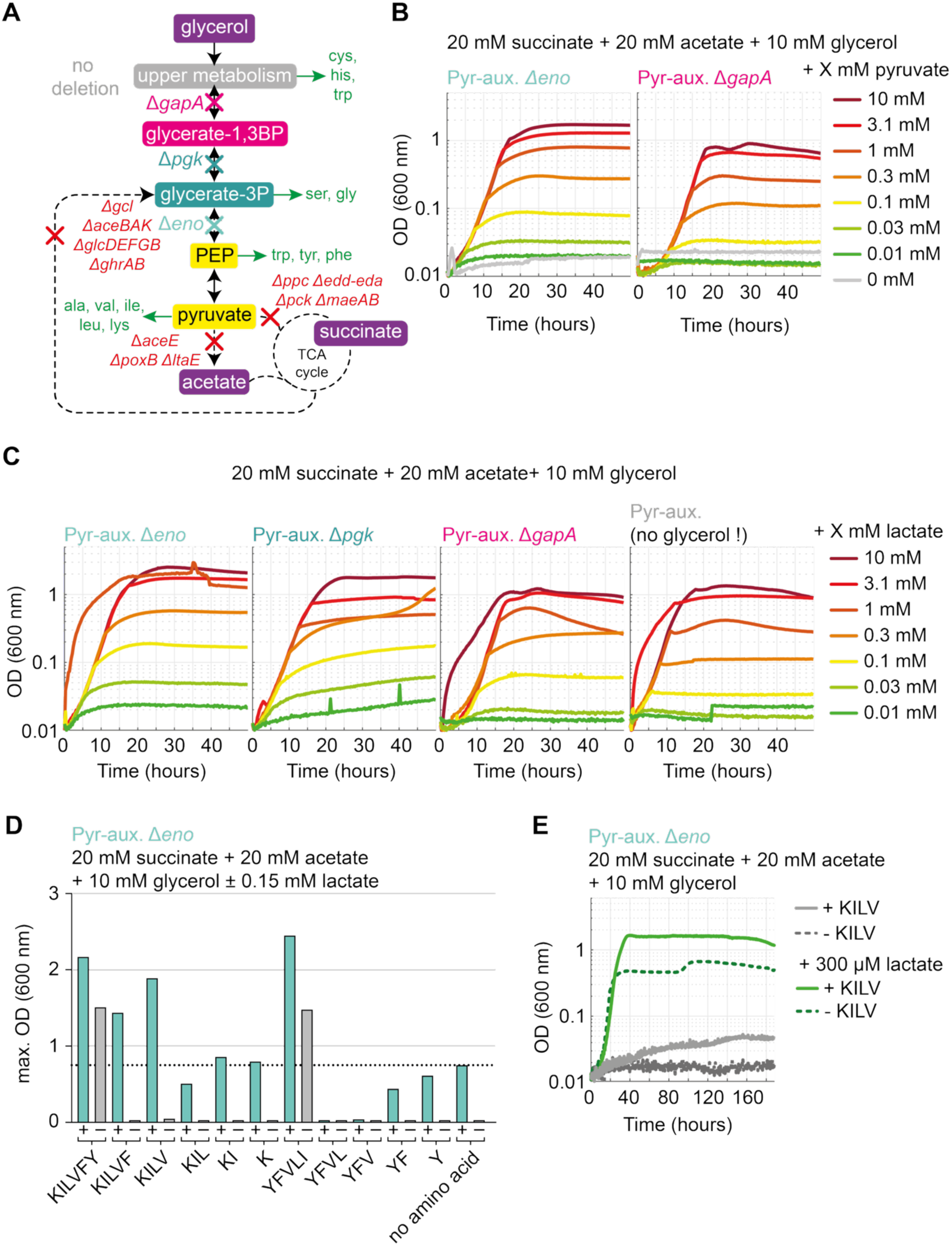
An array of pyruvate auxotrophic sensor strains. **A)** Selection scheme for pyruvate auxotroph strains. Pyruvate is separated from the TCA cycle by gene deletions (red) and needed as precursor for amino acids or upper metabolism (indicated in green). Strains with disrupted glycolysis are expected to have decreasing sensitivity with increasing number of biomass precursors which are selected for. Four different strains with increasing dependency on pyruvate and thus selective demand were created by deleting either *eno* (Pyr-aux. Δ*eno*), *pgk* (Pyr-aux. Δ*pgk*), *gapA* (Pyr-aux. Δ*gapA*) or leaving glycolysis open to replenish all upper metabolism from pyruvate (Pyr-aux.). **B)** Growth of Pyr-aux. Δ*eno* and Pyr-aux. Δ*gapA* with 10 mM glycerol + 20 mM acetate + 20 mM succinate and a gradient of pyruvate. **C)** Growth of all pyruvate sensors with 10 mM glycerol + 20 mM acetate + 20 mM succinate and a gradient of L-lactate. **D)** Maximal biomass yields (OD_600_) of Pyr-aux. Δ*eno* with amino acid supplementation. Growth on M9 + 20 mM succinate + 20 mM acetate + 10 mM glycerol with different combinations of amino acids (represented in one letter code, 1 mM each) with and without 0.15 mM L-lactate (indicated by + and – for each amino acid combination) was tested and compared to growth on M9 + 20 mM succinate + 20 mM acetate + 10 mM glycerol with 0.15 mM L-lactate (red bar to the right; final OD_600_ is indicated by red line). Values represent the mean values of two technical duplicates with error indicated by error bars. **E)** Growth of Pyr-aux. Δ*eno* with and without lactate supplementation with or without KILV (one letter amino acid code, 1 mM each) supplementation. Amino acid supplementation increases final OD_600_ in positive control media (dark and light green).

**Extended Data Figure 3:**
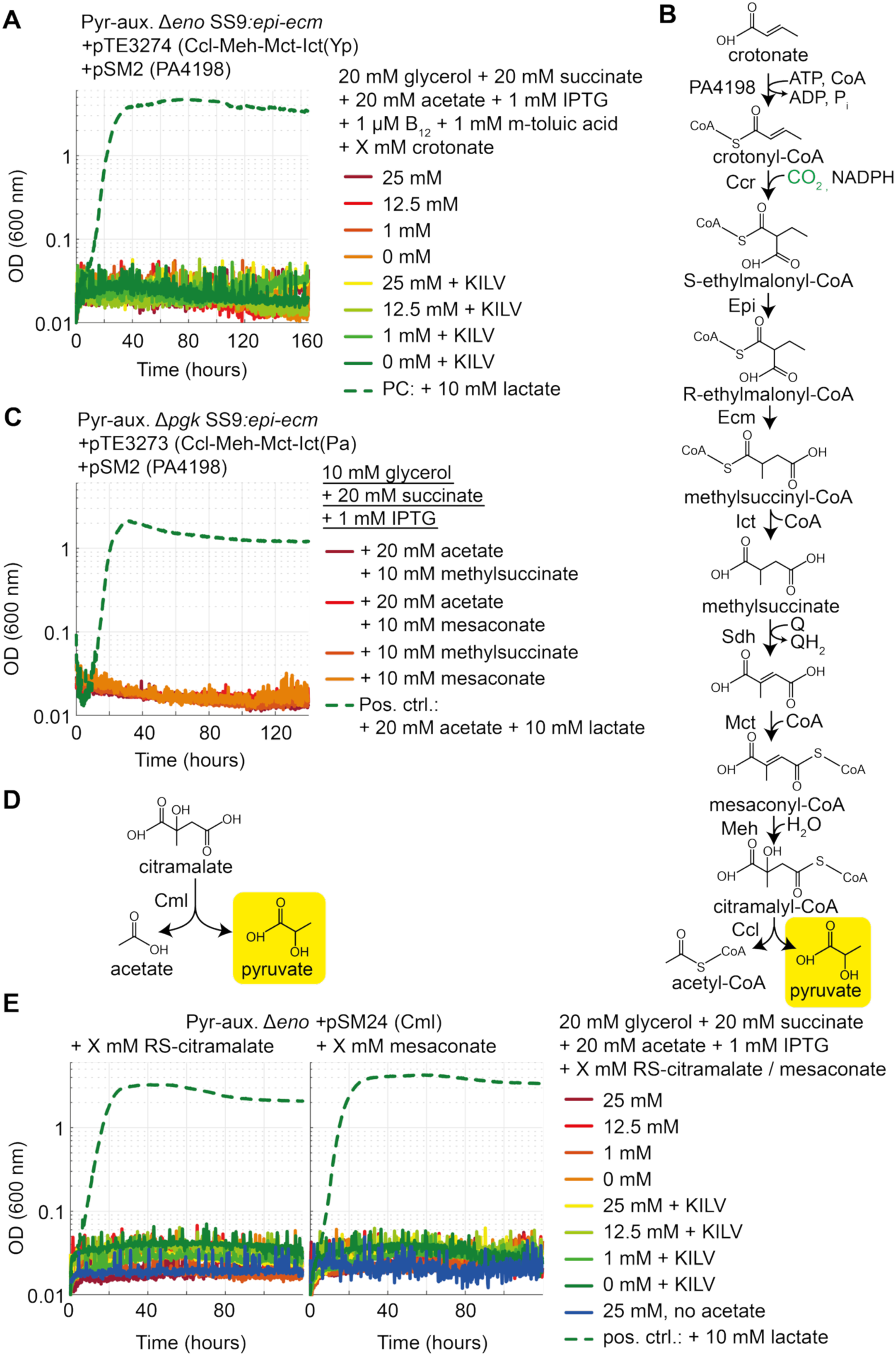
The pyruvate sensor strains do not grow via parts of the THETA cycle if mesaconate, methylsuccinate or crotonate are provided. **A)** The Pyr-aux. Δ*eno* does not grow in presence of crotonate if the expression of PA3198 is induced by addition of m-toluic acid. Growth is observed on positive control medium with lactate. **B)** Schematic representation of the reactions leading to pyruvate synthesis from crotonate with annotation of the corresponding enzymes. **C)** Growth of the Pyr-aux. Δ*pgk* expressing crotonate utilisation genes in presence of methylsuccinate, mesaconate, or lactate. Growth is only observed in presence of lactate (the positive control). To see if acetate supplementation was thermodynamically preventing Ccl dependent citramalyl-CoA cleavage, growth was also tested in absence of acetate. **D)** Schematic representation of the citramalate lyase activity of Cml from *Raoultella planticola.* **E)** The Pyr-aux. Δ*eno* strain expressing Cml does not grow with RS-citramalate with and without supplementation of the amino acids KILV (1 mM per amino acid). Growth is only observed on positive control medium with lactate.

**Extended Data Figure 4:**
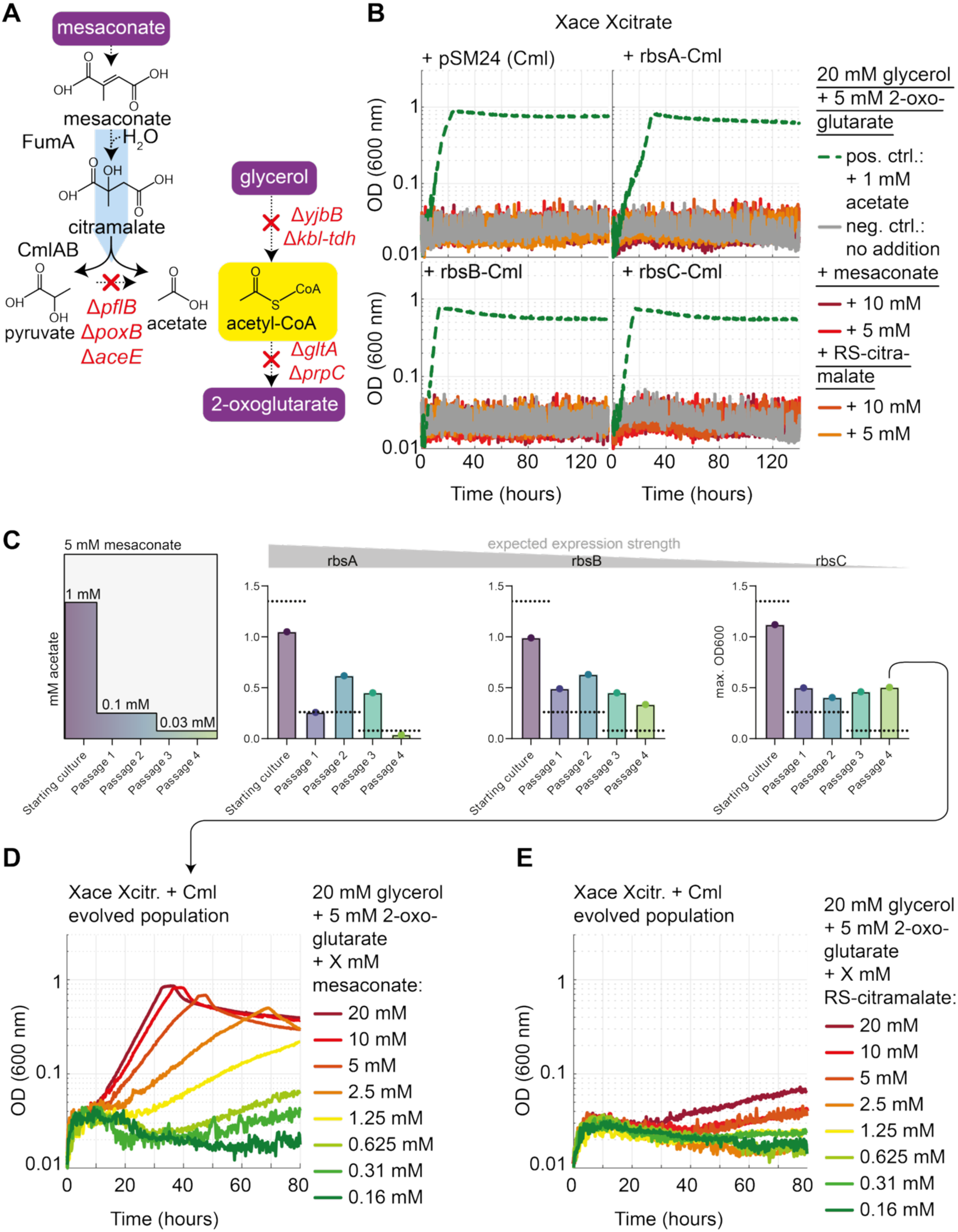
Evolution of mesaconate dependence. **A)** The endogenous Fumarase A (FumA) and a citramalate lyase (Cml) allow the conversion of mesaconate to pyruvate and acetate, which replenishes the metabolically isolated auxotrophy target acetyl-CoA. **B)** The unevolved Xace Xcitrate strain does not grow with mesaconate or RS-citramalate instead of acetate upon expressing Cml. Cml was expressed from pSM24 (previously tested in Pyr-aux. Δ*eno*, with a strong rbs^50^), or from pZ-ASS with rbsA (strong), rbsB (medium) or rbsC (weak)^28^. **C)** Semi-relaxing evolution for strains expressing Citramalate lyase with different expression levels. Dotted lines indicate the expected OD in case of pure acetate dependence. 20 mM glycerol + 5 mM 2-oxoglutarate were supplied. **D)** The evolved population expressing rbsC-*cmlAB* grows in a mesaconate dependent manner. **E)** The evolved population expressing rbsC-*cmlAB* does not grow with RS-citramalate instead of mesaconate.

**Extended Data Figure 5:**
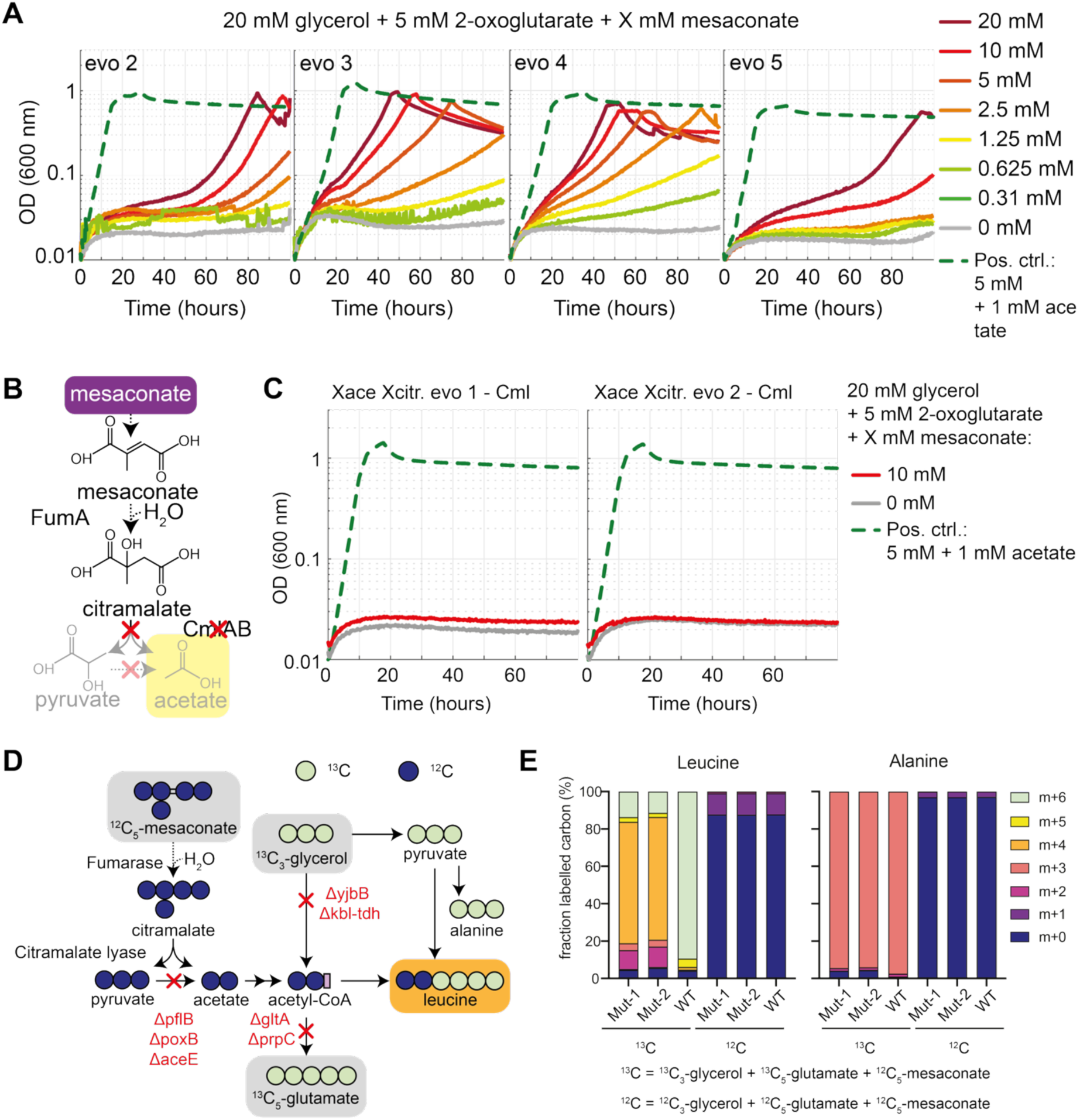
Characterisation of the mesaconate evolved isolates. **A)** Isolated single clones can grow with mesaconate. Clone 1 is shown in Figure 3. **B)** Removal of the citramalate lyase through plasmid curing should abolish growth of the Xace Xcitrate with mesaconate. **C)** The mesaconate evolved strains do not grow with mesaconate once cured of the citramalate lyase expression plasmid. **D)** Isotopic labelling patterns expected for mesaconate use via FumA and Cml. **E)** Leucine labelling confirms the desired incorporation of mesaconate derived carbon into acetyl-CoA.

**Extended Data Figure 6:**
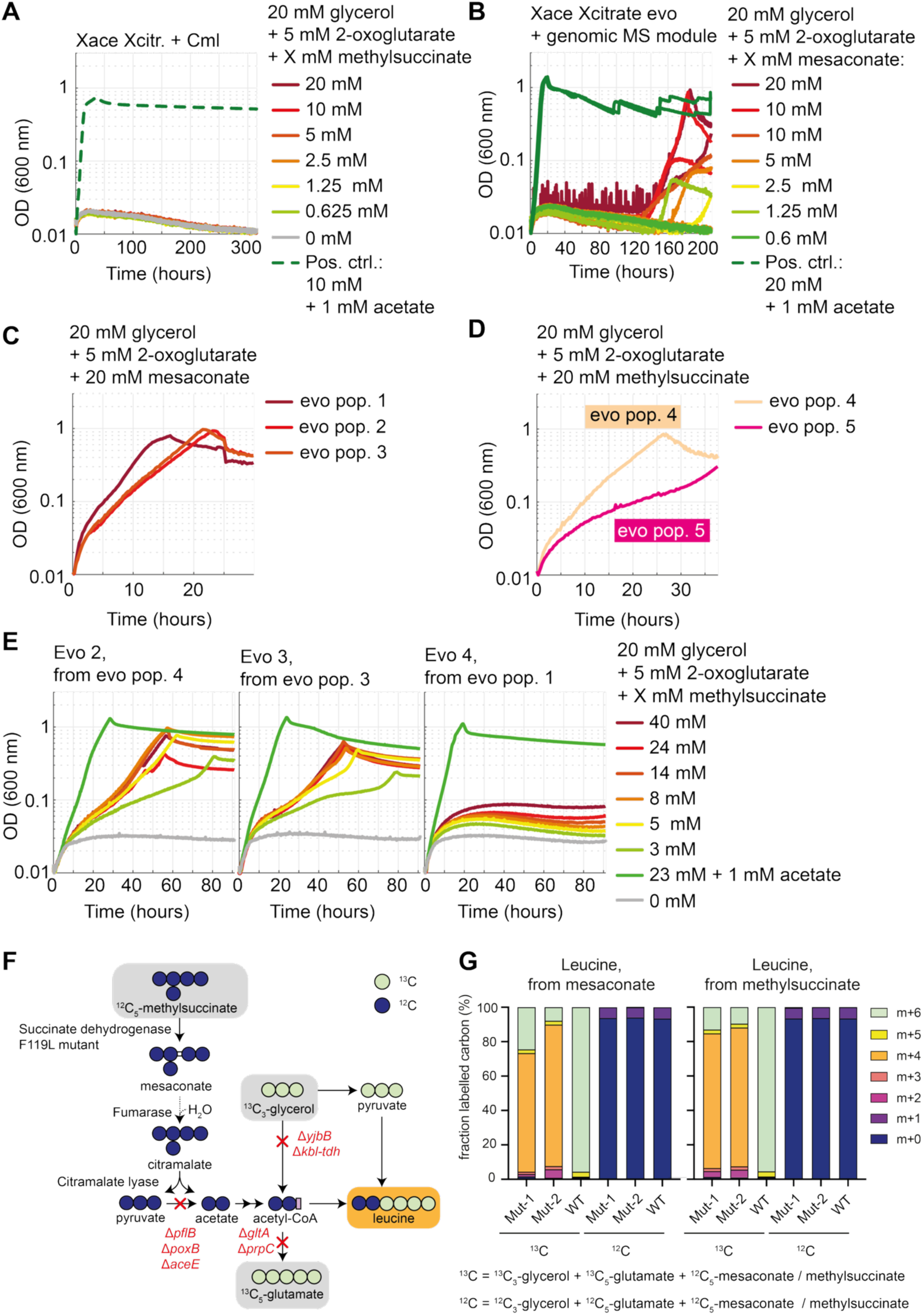
Characterisation of methylsuccinate evolved strains. **A)** The unevolved Xace Xcitrate strain expressing Cml cannot grow with methylsuccinate. **B)** Growth of the mesaconate evolved acetyl-CoA auxotroph with genomic integration of Cml and SdhA_F119L_SdhB in SS9. Populations evolve individually in both technical replicates to utilise the provided mesaconate after 120 hours. **C)** Growth of propagated populations capable of growing with mesaconate or **D)** methylsuccinate (populations 4 & 5). **E)** Growth of evolved clones 2-4 that were isolated from different populations with a gradient of methylsuccinate. Clone 4 was isolated from a population capable of growing with mesaconate, but not methylsuccinate. **F)** Isotopic labelling patterns expected for methylsuccinate use via Sdh and Cml. **G)** Leucine labelling confirms the desired incorporation of mesaconate and methylsuccinate in acetyl-CoA.

**Extended Data Figure 7:**
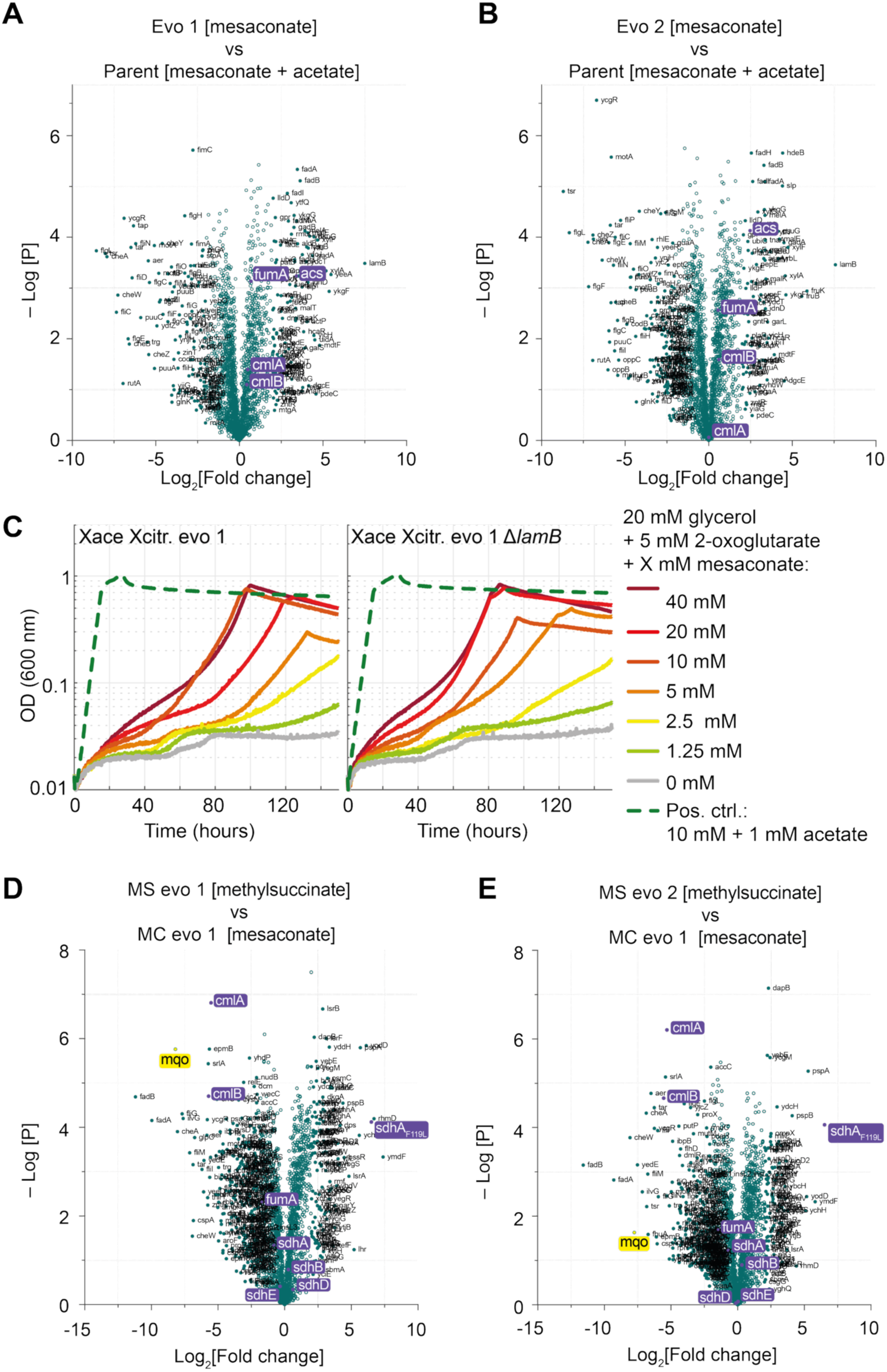
**A, B)** Proteomics comparison of mesaconate evolved isolates 1 (A) and 2 (B) grown with 20 mM glycerol + 10 mM mesaconate + 5 mM 2-oxoglutarate with the parental strain grown with 20 mM glycerol + 5 mM 2-oxoglutarate + 1 mM acetate. The pathway enzymes FumA and Acs are uregulated in both clones. **C)** Deleting *lamB* does not affect growth of the mesaconate evolved mutant 1 with mesaconate. Growth was tested on a gradient of mesaconate in presence of 20 mM glycerol + 5 mM 2-oxoglutarate. **D, E)** Proteomics comparison of the methylsuccinate evolved isolates 1 (E) and 2 (F) growing with methylsuccinate with its mesaconate evolved predecessor strain growin with mesaconate. 20 mM glycerol and 5 mM 2-oxoglutarate were supplied in both conditions.

**Extended Data Figure 8:**
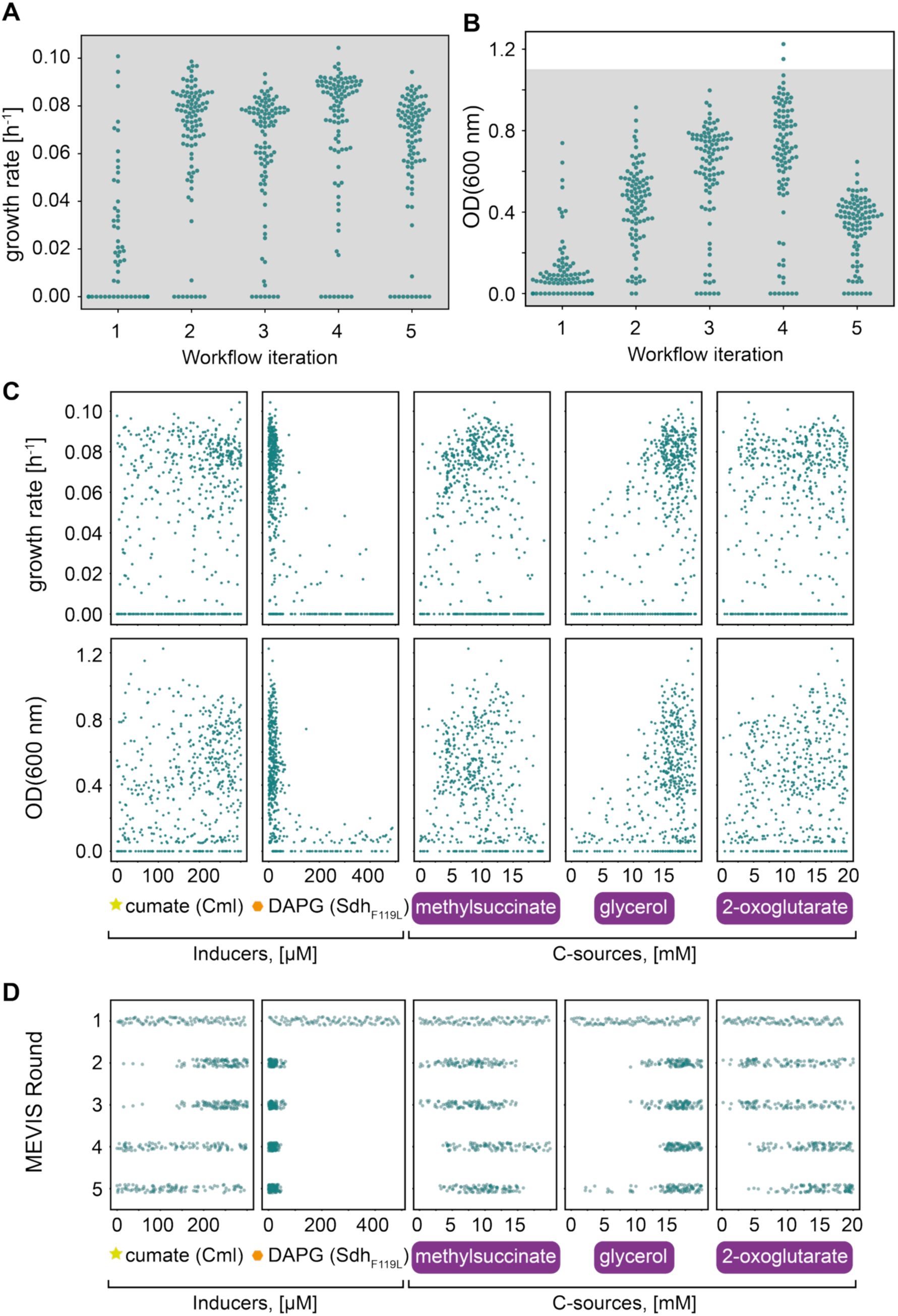
MEVIS for methylsuccinate dependent growth of an unevolved strain. **A)** Growth rates over five rounds of MEVIS. The grey threshold indicates the growth rate obtained through constitutive expression and adaptive laboratory evolution (0.11 h^-1^). **B)** Maximum optical density over five MEVIS rounds. The grey box indicates the OD of 1.1 obtained with the methylsuccinate evolved strain. **C)** Growth rates and maximal optical density of five MEVIS rounds over the tested concentrations of the inducers cumate and DAPG and carbon sources methylsuccinate, glycerol and 2-oxoglutarate. **D)** Tested concentrations for all components across MEVIS rounds 1-5.

**Extended Data Figure 9:**
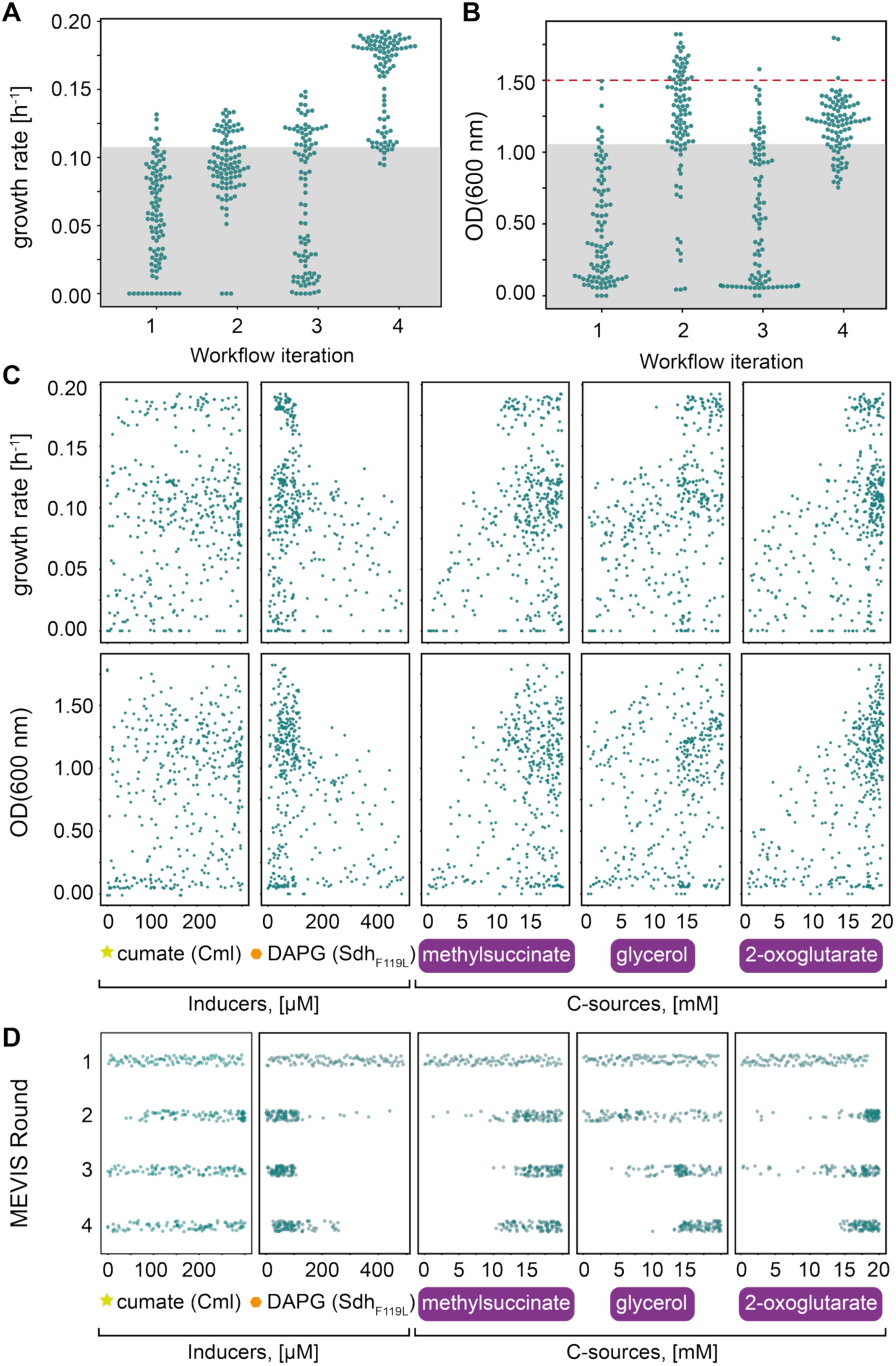
MEVIS for methylsuccinate dependent growth with the mesaconate evolved strain as background. **A)** Growth rates over four rounds of MEVIS. The grey threshold indicates the growth rate obtained through constitutive expression and adaptive laboratory evolution (0.11 h^-1^). **B)** Maximum optical density over four MEVIS rounds. The grey box indicates the OD of 1.1 obtained with the methylsuccinate evolved strain. The red line indicates the upper threshold ODs were constrained to when multiplying growth rates by OD to feed back to MEVIS, to enforce focusing on growth speed at high optical densities. **C)** Growth rates and maximal optical density of four MEVIS rounds over the tested concentrations of the inducers cumate and DAPG and carbon sources methylsuccinate, glycerol and 2-oxoglutarate. **C)** Tested concentrations for all components across MEVIS rounds 1-4.

