## Supplementary Information for "Combining evolution and machine learning-guided pathway optimization to engineer a novel methylsuccinate module for synthetic C1 metabolism *in vivo*"

[6] M4C-cluster of Excellence

### Supplementary Figures

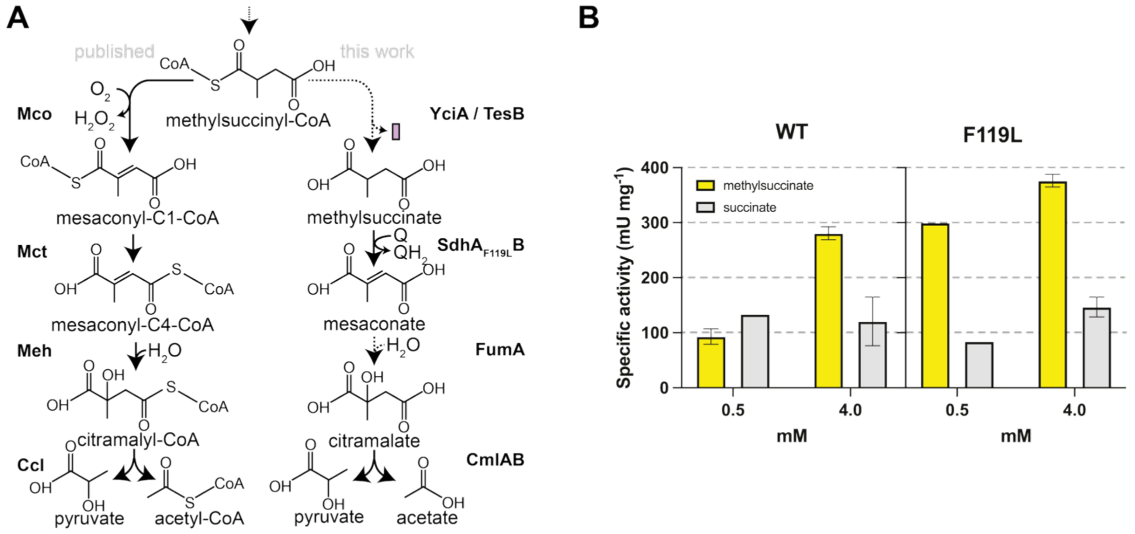


**Supplementary Figure 1: Redesign of the CoA ester-based THETA module towards an acid-based module variant. A)** Comparison of the published THETA module[1] for methylsuccinyl-CoA use and the module established in this work. **B)** Specific activities of wild type and F119L succinate dehydrogenase mutant with 0.5 mM and 4 mM of succinate or methylsuccinate. Ferrocenium reduction was followed by absorption measurements at 300 nm.

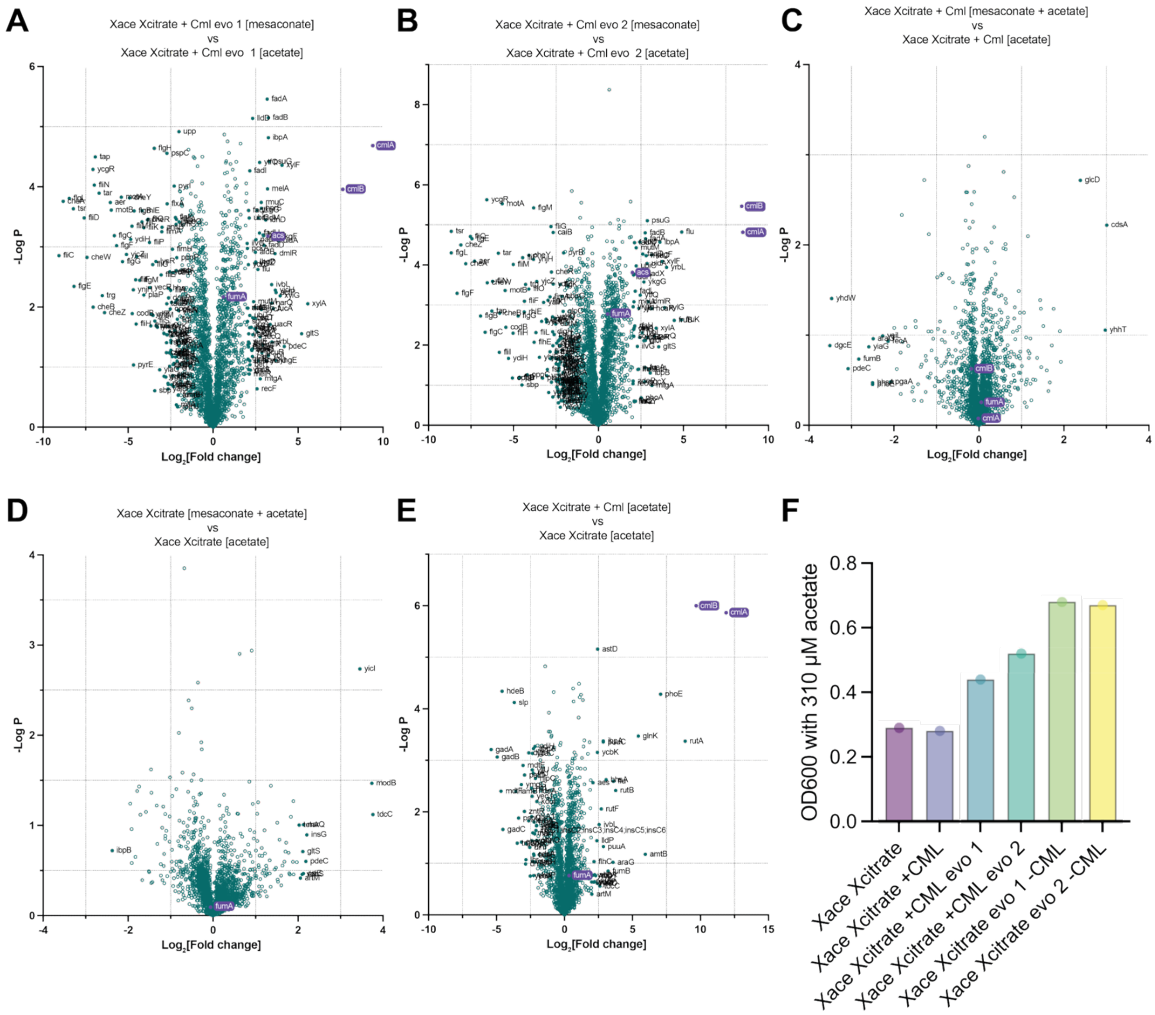


**Supplementary Figure 2: Proteomics comparisons of mesaconate utilising mutants.** Pathway enzymes are highlighted in violet. 20 mM glycerol + 5 mM 2-oxoglutarate were supplied in all conditions. **A)** Comparison of the mesaconate evolved clone 1 grown with mesaconate compared to acetate. **B)** Comparison of the mesaconate evolved clone 2 grown with mesaconate compared to acetate. **C)** Comparison of the parental acetyl-CoA auxotroph expressing Cml grown with mesaconate and acetate compared to acetate. **D)** Comparison of the parental acetyl-CoA auxotroph grown with mesaconate and acetate compared to acetate. **E)** Comparison of the parental acetyl-CoA auxotroph expressing Cml grown with acetate compared to the parental acetyl-CoA auxotroph grown with acetate. **F)** Optical densities achieved by the original Xace Xcitrate strain vs the evolved mutants with 310 µM acetate.

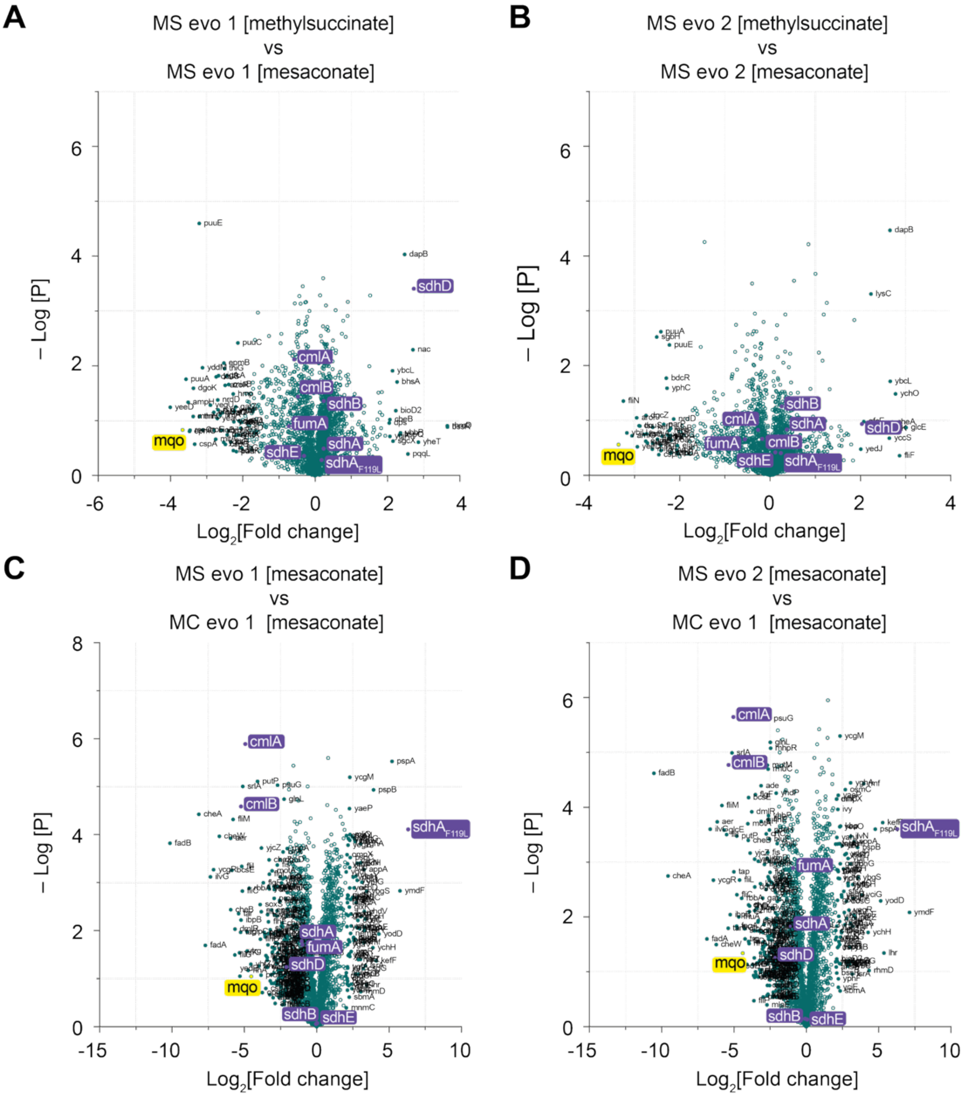


**Supplementary Figure 3: Proteomics comparisons for methylsuccinate evolved clones with mesaconate evolved clones.** 20 mM glycerol and 5 mM 2-oxoglutarate were provided in all conditions. Pathway enzymes are highlighted in violet. **A)** Comparison of the methylsuccinate evolved isolate 1 grown with methylsuccinate vs grown with mesaconate. **B)** Comparison of the methylsuccinate evolved isolate 2 grown with methylsuccinate vs grown with mesaconate. **C)** Comparison of the methylsuccinate evolved isolate 1 grown with mesaconate vs the mesaconate evolved predecessor strain grown with mesaconate. **D)** Comparison of the methylsuccinate evolved isolate 2 grown with mesaconate vs the mesaconate evolved predecessor strain grown with mesaconate.

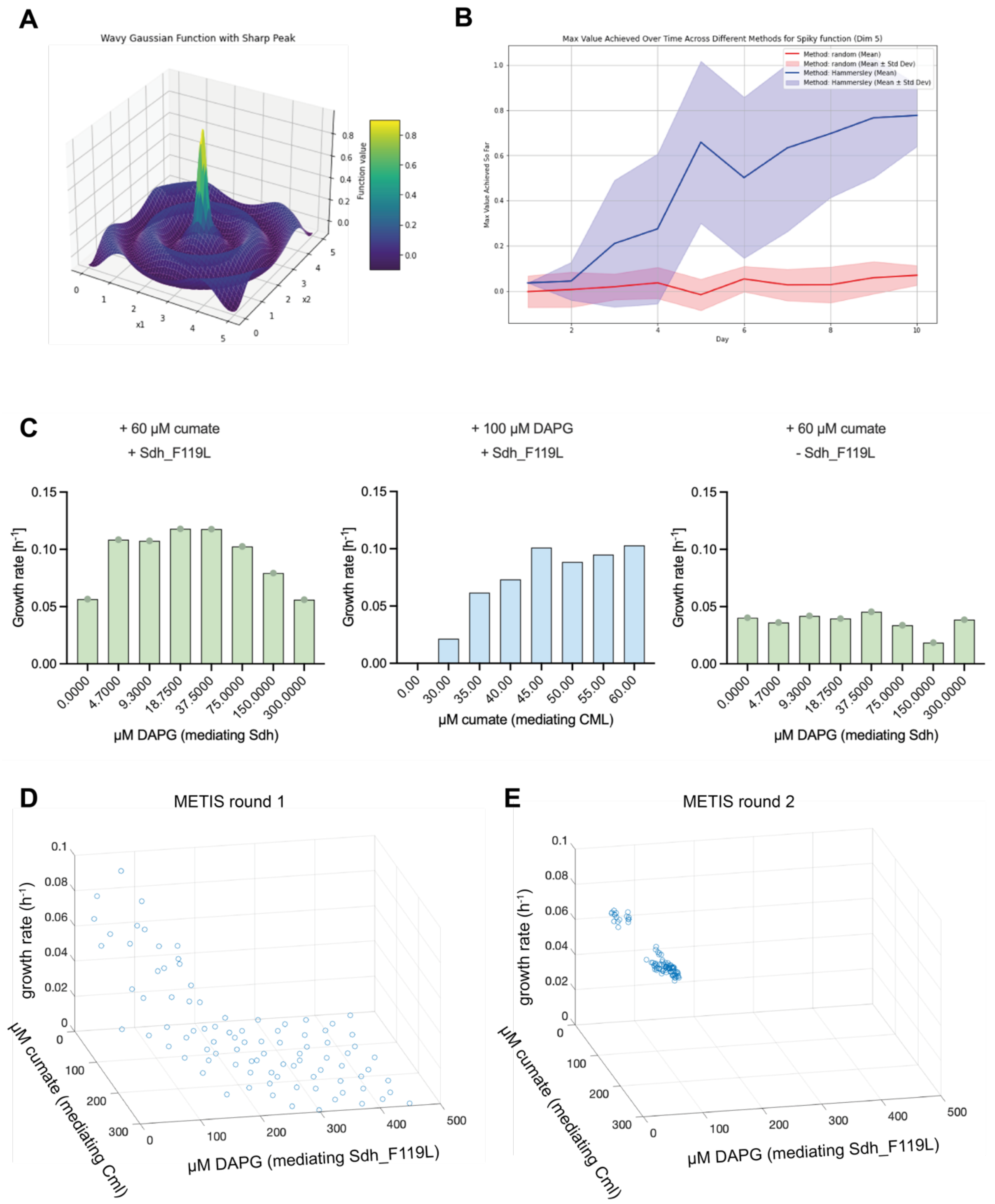


**Supplementary Figure 4: A)** Synthetic landscapes were used to assess the model performance. **B)** Hammersleigh sampling (blue) during the first day outperforms random sampling (red) for a three-dimensional landscape. **C)** Pre-screening results for inducible expressions of citramalate lyase and succinate dehydrogenase (F119L). Left graph: varying concentrations of DAPG, which mediates Sdh_F119L expression, while 60 µM cumate are supplied. Middle graph: varying concentrations of cumate, which mediates citramalate lyase expression, while 100 µM DAPG are supplied. Right: varying concentrations of DAPG in a strain only harbouring the native wild-type succinate dehydrogenase, while 60 µM cumate are supplied. **D)** Results of the first round of inducer combination tests proposed by METIS and pipetted using the ECHO liquid handling robot. The trends observed in pre-tests are confirmed. **E)** Results of the second round of inducer combination tests proposed by METIS based on the results of round 1.

### Supplementary Notes

#### Supplementary Note 1:

*Design of the MEVIS workflow for in vivo pathway engineering.*
To transfer the ability to fine-tune catalyst and substrate levels to the *in vivo* context in a workflow termed MEVIS (METIS for *in vivo* contexts), we improved the METIS[2] algorithm by changing the sampling method for the first screening round from random sampling to Hammersley sampling[3,4], thus testing concentrations dispersed more regularly for best solution landscape coverage, which resulted in faster predictions of global maxima during *in silico* testing (Supplementary Figure 11A, B). After adapting the METIS script for growth readouts, we aimed to optimize strain growth conditions by automated combinatorial pipetting with an Echo liquid handling robot and interpreting the data using growth rate as readout using METIS (Figure 4, MEVIS workflow). Here, the optimization of expression levels is based on the correlation between strain growth rate and inducer concentrations for rate-limiting pathway enzymes (Supplementary Figure 11C)[5]. We first applied the workflow to a combinatoric screen for the best expression levels for citramalate lyase and succinate dehydrogenase using growth rate as readout for pathway performance mwhile expressing both from two different, orthogonal inducible promoters from the Marionette strain collection[6] in the mesaconate evolved strains we had cured of the Cml plasmid (Supplementary Figure 11D, E). Here, reached an optimum of the solution landscape after one round of screening, as growth rates and yields showed little variation in round 2 (Supplementary Figure 11E).

**Supplementary Table 1: Genes upregulated more than 2-fold in both evolved mutants with mesaconate compared to the parental strain (with mesaconate and acetate) and the evolved mutants with mesaconate compared to the evolved mutants with acetate.** Information on cAMP dependent regulation was obtained from Ecocyc. Genes involved in acetate metabolism are highlighted in green, genes involved in lactate metabolism in orange.

| Gene | cAMP induced ? | Protein |
|---|---|---|
| *ykgF* | ? | putative amino acid dehydrogenase with NAD(P)-binding domain and ferridoxin-like domain |
| *xylA* | + | xylose isomerase |
| *xylF* | + | xylose ABC transporter periplasmic binding protein |
| *uidA* | + | β-D-glucuronidase |
| *psuG* | + | pseudouridine-5'-phosphate glycosidase |
| *yjcH* | + | DUF485 domain-containing inner membrane protein YjcH |
| *yrbL* | - | protein kinase-like domain-containing protein YrbL |

| | | |
|---|---|---|
| *ytfJ* | - | PF09695 family protein YtfJ |
| *melA* | + | α-galactosidase |
| *hcaR* | - | DNA-binding transcriptional dual regulator HcaR |
| *fadB* | - | multifunctional enoyl-CoA hydratase, 3-hydroxyacyl-CoA epimerase, $\Delta^3$-*cis*- $\Delta^2$-*trans*-enoyl-CoA isomerase, L-3-hydroxyacyl-CoA dehydrogenase |
| *xylG* | + | xylose ABC transporter ATP binding subunit |
| *fadA* | - | 3-ketoacyl-CoA thiolase |
| *acs* | + | acetyl-CoA synthetase (AMP-forming) |
| *fadH* | + | 2,4-dienoyl-CoA reductase |
| *ykgG* | ? | DUF162 domain-containing lactate utilization protein YkgG |
| *fadD* | + | long-chain-fatty-acid—CoA ligase |
| *rmuC* | - | putative recombination limiting protein RmuC |
| *ytfQ* | - | galactofuranose ABC transporter periplasmic binding protein |
| *ilvG* | ? | acetolactate synthase II subunit IlvG |
| *fucA* | + | L-fuculose-phosphate aldolase |
| *fadM* | - | long-chain acyl-CoA thioesterase FadM |
| *uacR* | ? | Putative σ-factor |
| *fadL* | + | long-chain fatty acid outer membrane channel / bacteriophage T2 receptor |
| *ubiC* | - | chorismate lyase |
| *lldD* | - | L-lactate dehydrogenase |

**Supplementary Table 1: Strains and plasmids used in this study.**

| **Strain** | **Strain, Genotype (Deletion)** |
|---|---|
| SL2 | YYC202 Δ*pflB* Δ*poxB* ΔaceA Δ*kbl* Δ*ltaE* |
| Xace Xcitrate | SIJ488 Δ*pflB*, Δ*poxB*, Δ*aceE*, Δ*gltA*, Δ*prpC*, Δ*yjbB*,Δ*kbl-tdh* |
| Xace Xcitrate + CML | SIJ488 Δ*pflB*, Δ*poxB*, Δ*aceE*, Δ*gltA*, Δ*prpC*, Δ*yjbB*,Δ*kbl-tdh* pZ-ASS-rbsC-*cmlAB* |
| Xace Xcitrate + CML evolved population | SIJ488 Δ*pflB*, Δ*poxB*, Δ*aceE*, Δ*gltA*, Δ*prpC*, Δ*yjbB*,Δ*kbl-tdh* pZ-ASS-rbsC-*cmlAB*, mesaconate evolved mixed population |
| Xace Xcitrate + CML evo 1 | SIJ488 Δ*pflB*, Δ*poxB*, Δ*aceE*, Δ*gltA*, Δ*prpC*, Δ*yjbB*,Δ*kbl-tdh* pZ-ASS-rbsC-*cmlAB*, mesaconate evolved single clone isolate 1 |
| Xace Xcitrate + CML evo 2 | SIJ488 Δ*pflB*, Δ*poxB*, Δ*aceE*, Δ*gltA*, Δ*prpC*, Δ*yjbB*,Δ*kbl-tdh* pZ-ASS-rbsC-*cmlAB*, mesaconate evolved single clone isolate 2 |
| Xace Xcitrate evo 1 cured | SIJ488 Δ*pflB*, Δ*poxB*, Δ*aceE*, Δ*gltA*, Δ*prpC*, Δ*yjbB*,Δ*kbl-tdh* mesaconate evolved single clone isolate 1 |
| Xace Xcitrate evo 2 cured | SIJ488 Δ*pflB*, Δ*poxB*, Δ*aceE*, Δ*gltA*, Δ*prpC*, Δ*yjbB*,Δ*kbl-tdh* mesaconate evolved single clone isolate 2 |
| Xace Xcitrate + CML Sdh$_{mut}$ | SIJ488 Δ*pflB*, Δ*poxB*, Δ*aceE*, Δ*gltA*, Δ*prpC*, Δ*yjbB*,Δ*kbl-tdh* Δ*fadB* SS7-*epi-ecm* SS9-rbsC-*cmlAB*-rbsC-*sdhA*_F119L-*sdhB* |
| Xace Xcitrate + CML Sdh$_{mut}$ evo pop. 4 | SIJ488 Δ*pflB*, Δ*poxB*, Δ*aceE*, Δ*gltA*, Δ*prpC*, Δ*yjbB*,Δ*kbl-tdh* Δ*fadB* SS7-*epi-ecm* SS9-rbsC-*cmlAB*-rbsC-*sdhA*_F119L-*sdhB*, methylsuccinate evolved mixed population |

| | |
|---|---|
| Xace Xcitrate + CML $Sdh_{mut}$ evo pop. 5 | SIJ488 Δ*pflB*, Δ*poxB*, Δ*aceE*, Δ*gltA*, Δ*prpC*, Δ*yjbB*,Δ*kbl-tdh* Δ*fadB* SS7-*epi-ecm* SS9-rbsC-*cmlAB*-rbsC-s*dhA*_F119L-*sdhB*, methylsuccinate evolved mixed population |
| Xace Xcitrate + CML $Sdh_{mut}$ evo 1 | SIJ488 Δ*pflB*, Δ*poxB*, Δ*aceE*, Δ*gltA*, Δ*prpC*, Δ*yjbB*,Δ*kbl-tdh* Δ*fadB* SS7-*epi-ecm* SS9-rbsC-*cmlAB*-rbsC-s*dhA*_F119L-*sdhB*, methylsuccinate evolved single clone isolate 1 |
| Xace Xcitrate + CML $Sdh_{mut}$ evo 2 | SIJ488 Δ*pflB*, Δ*poxB*, Δ*aceE*, Δ*gltA*, Δ*prpC*, Δ*yjbB*,Δ*kbl-tdh* Δ*fadB* SS7-*epi-ecm* SS9-rbsC-*cmlAB*-rbsC-s*dhA*_F119L-*sdhB*, methylsuccinate evolved single clone isolate 2 |
| Xace Xcitrate + CML $Sdh_{mut}$ evo 3 | SIJ488 Δ*pflB*, Δ*poxB*, Δ*aceE*, Δ*gltA*, Δ*prpC*, Δ*yjbB*,Δ*kbl-tdh* Δ*fadB* SS7-*epi-ecm* SS9-rbsC-*cmlAB*-rbsC-s*dhA*_F119L-*sdhB*, methylsuccinate evolved single clone isolate 3 |
| Xace Xcitrate + CML $Sdh_{mut}$ evo 4 | SIJ488 Δ*pflB*, Δ*poxB*, Δ*aceE*, Δ*gltA*, Δ*prpC*, Δ*yjbB*,Δ*kbl-tdh* Δ*fadB* SS7-*epi-ecm* SS9-rbsC-*cmlAB*-rbsC-s*dhA*_F119L-*sdhB*, methylsuccinate evolved single clone isolate 4 |
| Xace Xcitrate + crotonate | SIJ488 Δ*pflB*, Δ*poxB*, Δ*aceE*, Δ*gltA*, Δ*prpC*, Δ*yjbB*,Δ*kbl-tdh* Δ*fadB* SS7-*epi-ecm* SS9-rbsC-*cmlAB*-rbsC-s*dhA*_F119L-*sdhB*, methylsuccinate evolved single clone isolate 1 pZ-ASS-DmdB1-Ccr |
| Xace Xcitrate + crotonate FumA | SIJ488 Δ*pflB*, Δ*poxB*, Δ*aceE*, Δ*gltA*, Δ*prpC*, Δ*yjbB*,Δ*kbl-tdh* Δ*fadB* SS7-*epi-ecm* SS9-rbsC-*cmlAB*-rbsC-s*dhA*_F119L-*sdhB*, methylsuccinate evolved single clone isolate 1 pZ-ASS-DmdB1-Ccr-FumA |
| Xace Xcitrate + crotonate FumA TesB | SIJ488 Δ*pflB*, Δ*poxB*, Δ*aceE*, Δ*gltA*, Δ*prpC*, Δ*yjbB*,Δ*kbl-tdh* Δ*fadB* SS7-*epi-ecm* SS9-rbsC-*cmlAB*-rbsC-s*dhA*_F119L-*sdhB*, methylsuccinate evolved single clone isolate 1 pZ-ASS-DmdB1-Ccr-FumA-TesB |
| Xace Xcitrate + crotonate FumA YciA | SIJ488 Δ*pflB*, Δ*poxB*, Δ*aceE*, Δ*gltA*, Δ*prpC*, Δ*yjbB*,Δ*kbl-tdh* Δ*fadB* SS7-*epi-ecm* SS9-rbsC-*cmlAB*-rbsC-s*dhA*_F119L-*sdhB*, methylsuccinate evolved single clone isolate 1 pZ-ASS-DmdB1-Ccr-FumA-YciA |
| Xace Xcitrate + Mar-Cml | SIJ488 Δ*pflB*, Δ*poxB*, Δ*aceE*, Δ*gltA*, Δ*prpC*, Δ*yjbB*,Δ*kbl-tdh* mesaconate evolved single clone isolate 1 (Xace Xcitrate evo 1 cured) +pAJM.657-rbsC-*cmlAB* |
| Xace Xcitrate + Mar-Cml Mar-Sdh | SIJ488 Δ*pflB*, Δ*poxB*, Δ*aceE*, Δ*gltA*, Δ*prpC*, Δ*yjbB*, Δ*kbl-tdh* mesaconate evolved single clone isolate 1 (Xace Xcitrate evo 1 cured) +pAJM.657-rbsC-*cmlAB* +pSEVA624-$P_{PhlF}$-s*dhA*_F119L-*sdhB* _PhlFAM |
| Xace Xcitrate +GltA | SIJ488 Δ*pflB*, Δ*poxB*, Δ*aceE*, Δ*gltA*, Δ*prpC*, Δ*yjbB*, Δ*kbl-tdh* +pSEVA-224-GltA |
| Pyr-aux. | SIJ488 Δ*gcl* Δ*aceBAK* Δ*glcDEFGB* Δ*ghrB* Δ*ghrA* Δ*ppc* Δ*maeA* Δ*pck* Δ*maeB* Δ*edd-eda* Δ*aceE*::Kan |
| Pyr-aux. Δ*pgk* | SIJ488 Δ*gcl* Δ*aceBAK* Δ*glcDEFGB* Δ*ghrB* Δ*ghrA* Δ*ppc* Δ*maeA* Δ*pck* Δ*maeB* Δ*edd-eda* Δ*aceE* Δ*pgk* SS9::rbsB::*epi-ecm* |
| Pyr-aux. Δ*eno* | SIJ488 Δ*gcl* Δ*aceBAK* Δ*glcDEFGB* Δ*ghrB* Δ*ghrA* Δ*ppc* Δ*maeA* Δ*pck* Δ*maeB* Δ*edd-eda* Δ*aceE* Δ*mgsA* Δ*yciA* Δ*tesB* Δ*poxB-ltaE* Δ*eno*::CapR SS9::rbsB::*epi-ecm* |
| Pyr-aux. Δ*gapA* | SIJ488 Δ*gcl* Δ*aceBAK* Δ*glcDEFGB* Δ*ghrB* Δ*ghrA* Δ*ppc* Δ*maeA* Δ*pck* Δ*maeB* Δ*edd-eda* Δ*aceE* Δ*mgsA* Δ*yciA* Δ*tesB* Δ*poxB-ltaE* Δ*gapA*::CapR SS9::rbsB::*epi-ecm* |
| Pyr-aux. Δ*pgk* +pTE3273 +pSM2 | SIJ488 Δ*gcl* Δ*aceBAK* Δ*glcDEFGB* Δ*ghrB* Δ*ghrA* Δ*ppc* Δ*maeA* Δ*pck* Δ*maeB* Δ*edd-eda* Δ*aceE* Δ*pgk* SS9::rbsB::*epi-ecm* + pTE3273 (pLlacO1-Ccl-Meh-Mct-Ict (Pa) (no LacI)) +pSM2 (Pm-PA4198) |
| Pyr-aux. Δ*eno* +pTE3274 +pSM2 | SIJ488 Δ*gcl* Δ*aceBAK* Δ*glcDEFGB* Δ*ghrB* Δ*ghrA* Δ*ppc* Δ*maeA* Δ*pck* Δ*maeB* Δ*edd-eda* Δ*aceE* Δ*mgsA* Δ*yciA* Δ*tesB* Δ*poxB-ltaE* Δ*eno*::CapR SS9::rbsB::*epi-ecm* + pTE3273 (pLlacO1-Ccl-Meh-Mct-Ict (Yp) (no LacI)) +pSM2 (Pm-PA4198) |

| | |
|---|---|
| Pyr-aux. Δ*eno* +pSM24 | SIJ488 Δ*gcl* Δ*aceBAK* Δ*glcDEFGB* Δ*ghrB* Δ*ghrA* Δ*ppc* Δ*maeA* Δ*pck* Δ*maeB* Δ*edd-eda* Δ*aceE* Δ*mgsA* Δ*yciA* Δ*tesB* Δ*poxB-ltaE* Δ*eno*::CapR SS9::rbsB::*epi-ecm* +pSM24 (pLlacO1-Cml) |
| Succinate auxotroph | SIJ488 Δ*gcl* Δ*aceBAK* Δ*glcGB* Δ*gltA* Δ*sucAB* Δ*ghrB* Δ*ghrA* Δ*prpC* Δ*sdhABCD* Δ*frdABCD* SS9:aceA:KanR |
| Succinate auxotroph +Ccr +MeaB-Epi-Mcm | SIJ488 Δ*gcl* Δ*aceBAK* Δ*glcGB* Δ*gltA* Δ*sucAB* Δ*ghrB* Δ*ghrA* Δ*prpC* Δ*sdhABCD* Δ*frdABCD* SS9:aceA:KanR +pTE3288 (pLlacO1-DamdB1-Ccr) +pSM183 (PPhlF-MeaB-Epi-Ecm) |
| 2-OXO-AUX[7] | Δ*gcl* Δ*aceBAK* Δ*glcGB* Δ*gltA* Δ*sucAB* Δ*ghrB* Δ*ghrA* Δ*prpC* SS9:*aceA*:KanR |
| 2-OXO-AUX SS7:*epi-ecm* | Δ*gcl* Δ*aceBAK* Δ*glcGB* Δ*gltA* Δ*sucAB* Δ*ghrB* Δ*ghrA* Δ*prpC* SS9:*aceA*:KanR SS7:*epi-ecm* |
| 2-OXO-AUX SS7:*epi-ecm* +pSM7 | Δ*gcl* Δ*aceBAK* Δ*glcGB* Δ*gltA* Δ*sucAB* Δ*ghrB* Δ*ghrA* Δ*prpC* SS9:*aceA*:KanR SS7:*epi-ecm* +pSM7 (Ptrc-Mch-Mcl) |
| C1+GLYAUX[7] | Δ*gcl* Δ*aceBAK* Δ*glcDEFGB* Δ*glyA* Δ*ltaE* Δ*ghrB* Δ*ghrA* Δ*kbl-tdh* |
| C1+GLYAUX SS7:*epi-ecm* | Δ*gcl* Δ*aceBAK* Δ*glcDEFGB* Δ*glyA* Δ*ltaE* Δ*ghrB* Δ*ghrA* Δ*kbl-tdh* SS7:*epi-ecm* |
| C1+GLYAUX SS7:*epi-ecm* +pSM7 | Δ*gcl* Δ*aceBAK* Δ*glcDEFGB* Δ*glyA* Δ*ltaE* Δ*ghrB* Δ*ghrA* Δ*kbl-tdh* SS7:*epi-ecm* +pSM7 (Ptrc-Mch-Mcl) |
| TCA-AUX[7] | Δ*gcl* Δ*aceBAK* Δ*glcDEFGB* Δ*ghrB* Δ*ghrA* Δ*ppc* Δ*maeA* Δ*pck* Δ*maeB* SS9:*glcB* |
| TCA-AUX +pSM7 | Δ*gcl* Δ*aceBAK* Δ*glcDEFGB* Δ*ghrB* Δ*ghrA* Δ*ppc* Δ*maeA* Δ*pck* Δ*maeB* SS9:*glcB* +pSM7 (Ptrc-Mch-Mcl) |

**Supplementary Table 3. Oligonucleotide primers used**. 'KO' primers were used for amplification of the CapR cassette from pKD3 (overlap with CapR sequence marked in bold) with 50 bp overhangs homologous to regions upstream and downstream of the target locus (Purpose 1). 'KO_Ver'-primers were used to verify successful by CapR resistance cassette insertion as well as flippase mediated cassette removal (Purpose 2). Cloning primers are marked with Purpose 3.

| name | Sequence (5' → 3')) | purpose |
|---|---|---|
| *lldD*_KO_ F | CCCGCCTGCCCGGTGAGCATAATGAGCATTCGAGGGAGAAAAACGCATGA**GAGTAGGGAACTGCCAGGCA** | 1 |
| *lldD*_KO_R | AGAGGGTTAGGGTGAGGGGGCGCAAACGACTATGCCGCATTCCCTTTCGC**GGTCCATATGAATATCCTCCTTAGTTCCT** | 1 |
| *lldD*_KO_Ver_F | CATGCGCGGTTTCTTCGATGTC | 2 |
| *lldD*_KO_Ver_R | TGCGGTGTCGTTTCAGAGTGAG | 2 |
| *aceBAK*_KO_F | TCGTTCACAGTGGGGAAGTTTTCGGATCCATGACGAGGAGCTGCACGATG**GTGTAGGCTGGAGCTGCTTC** | 1 |
| *aceBAK*_KO_R | TGCGGAGAAAAATTATATGGAAGCTTTACTCAAAAAAGCATCTCCCCATA**GGAATTAGCCATGGTCCATATG** | 1 |
| *aceB*_KO_Ver_F | TCCGAAACGTACCTCAGCAG | 2 |
| *aceK*_KO_Ver_R | GCTTTATCTGACCATGCACGC | 2 |
| *glcDEF*_KO_F | GAATGACTTTAGTTTTCATTTTGTTATTCCTTTTCAAGGGCTTGTTCTAC**GTGTAGGCTGGAGCTGCTTC** | 1 |
| *glcDEF*_KO_R | CGCGCAAAAATCAGCTGCCACACAACACAACAAAGCGAAGCCTACTCATG**GGAATTAGCCATGGTCCATATG** | 1 |
| *glcDEF*_KO_Ver_F | CGCAGTCACGAACGCGGTACGT | 2 |
| *glcDEF*_Ver_R | CACACAGTCGACGTTCCGAGGGAAG | 2 |
| *ghrB*_KO_F | CATATTTCAGGCTAAGGTGATCGCCTTATCAGTGAATGGAGAGAAGCATG**GTGTAGGCTGGAGCTGCTTC** | 1 |
| *ghrB*_KO_R | TATCGGGCTTTACTCTACGCAGTCGCGGCTTAGTCCGCGACGTGCGGATT**GGAATTAGCCATGGTCCATATG** | 1 |
| *ghrB*_KO_Ver_F | GGGCCTTGGCCCGGCTAACC | 2 |
| *ghrB*_KO_Ver_R | GCTTTGTCGCAAATGGCATC | 2 |
| *ghrA*_KO_F | AACGATAAGTGCGAATAAATTTCGCACAACGCTTTTCGGGAGTCAGTATG**GTGTAGGCTGGAGCTGCTTC** | 1 |

| | | |
|---|---|---|
| *ghrA*_KO_R | CCAAGGATAGCAGGAATCCTGATGCTTTATTAGTAGCCGCGTGCGCGGTC**GGAATTAGCCATGGTCCATATG** | 1 |
| *ghrA*_KO_Ver_F | CTGGAACGGGCGCTAATTTAG | 2 |
| *ghrA*_KO_Ver_R | GGGCCATGATCGGTGATCGC | 2 |
| *ppc*_KO_F | AAAGCACGAGGGTTTGCAGAAGAGGAAGATTAGCCGGTATTACGCATACC**GTGTAGGCTGGAGCTGCTTC** | 1 |
| *ppc*_KO_R | CAAACGATAAGATGGGGTGTCTGGGGTAATATGAACGAACAATATTCCGC**GGAATTAGCCATGGTCCATATG** | 1 |
| *ppc*_KO_Ver_F | ACGAGGGTGTTAGAACAGAAGT | 2 |
| *ppc*_KO_Ver_R | CAAAGCCCGAGCATATTCGC | 2 |
| *maeB*_KO_F | TTCAGGGTAAGCGTGAGAGTTAAAAAAAATTACAGCGGTTGGGTTTGCGC**GTGTAGGCTGGAGCTGCTTC** | 1 |
| *maeB*_KO_R | TTGCCCACACACTTTATTTGTGAACGTTACGTGAAAGGAACAACCAAATG**GGAATTAGCCATGGTCCATATG** | 1 |
| *maeB*_KO_Ver_F | AGAGATATTCGCTGTGGTGCA | 2 |
| *maeB*_KO_Ver_R | GCAGACAGGCATGGTATTGC | 2 |
| *eda*_KO_F | GCCCGATCCTCGATCGGGCATTTTGACTTTTACAGCTTAGCGCCTTCTAC**GTGTAGGCTGGAGCTGCTTC** | 1 |
| *edd*_KO_R | CGCGTTGTGAATCATCCTGCTCTGACAACTCAATTTCAGGAGCCTTTATG**GGAATTAGCCATGGTCCATATG** | 1 |
| *eda*_KO_Ver_F | GGAAATTGATGGCAGCCGTC | 2 |
| *edd*_KO_Ver_R | CCGGTTACAGGCGTTTCAGTCA | 2 |
| *yciA*_KO_F | TCATAGTAGCATCGCGCCTGTGATTTTCCTTTTAAGTCGGTTTTACCATG**GTGTAGGCTGGAGCTGCTTC** | 1 |
| *yciA*_KO_R | AAAAGCCTCCGACCGGAGGCTTTTGACTATTACTCAACAGGTAAGGCGCG**GGAATTAGCCATGGTCCATATG** | 1 |
| *yciA*_KO_Ver_F | ACATCGCATTCTGGCTGC | 2 |
| *yciA*_KO_Ver_R | CGTAATCAGCCGCAGTATCC | 2 |
| *ltaE*_KO_F | CTAAAATGCGTTGCGGCACGTCTCTCTCCTTAACGCGCCAGGAATGCACG**GTGTAGGCTGGAGCTGCTTC** | 1 |
| *poxB*_KO_R | GATGAACTAAACTTGTTACCGTTATCACATTCAGGAGATGGAGAACCATG**GGAATTAGCCATGGTCCATATG** | 1 |
| *ltaE*_KO_Ver_R | AGGATCTGATGCCCTTGCTG | 2 |
| *poxB*_KO_Ver_F | AGTGCCTCCTTTCTCTCCCA | 2 |
| *tesB*_KO_Ver_F | CAATCGCAACCGCTAAACC | 2 |
| *tesB*_KO_Ver_R | TTCGCTGAAGGTGTGACG | 2 |
| pZ-ASS-F | GCATTTATCAGGGTTATTGTCTCATG | 3 |
| pZ-ASS-R | CTAGGGCGGCGGATTTGTCCTAC | 3 |
| pKD3-F | ATAATACCGCGCCACATAGC | 3 |
| pKD3-R | CATATGAATATCCTCCTTAGTTCCT | 3 |
| pKD3-in-SS9-F | GGAGAGCGTTTTCAATCCTACCTCTGGCGCAGTTGATATGTAAGGCAGGTGAGAGTAGGGAACTGCCAGG | 3 |
| pKD3-in-SS9-R | AATTGCCGTTAAAACTAAAAACAGCATCAATAATCAACGCGATATAATAAATGGGAATTAGCCATGGTCC | 3 |
| Ec162_poxB_ver_R | Catggcgcggcttggtcgggtaacggtatcactgcgtaaatcaatcatgg | 2 |
| Ec161_poxB_ver_F | Agcaggccagtacgtaaccttcagcgattcggcgtcggtcagcgtcatt | 2 |
| Ec104_kbl-tdh_ver_R | Gtgcggaaatcatcgcccgcatttatataacggaataatacactaagttt | 2 |
| Ec103_kbl-tdh_ver_F | Tatcaaaacgctacccactaatggagggaaacctattttcttgaaaggaa | 2 |
| Ec253_yjbB_V_R | Gcttcttcaccagccttaaacacataagtggtgttcgggccacccagcgtagcgtgacct | 2 |
| Ec252_yjbB_V_F | Accgaactggggtatgacgcggactgattcacaaatcgtcacttttcc | 2 |
| Ec249_pflB_V_R | Gttccacaggattcaaaggagtgaatgcgaccaataactgacattgcggt | 2 |
| MR937_gltA_Ver_R | Ctctgtgaaagagggaaaacctgggtacagagctctgggcgcttgcagg | 2 |
| MR936_gltA_Ver_F | Aattgaaagtattgggtgctgataattgagctgtctattctttttaa | 2 |
| MR935_prpC_Ver_R | tacgaataacaataaggaaacgtacccaagtcagctcaaatcaacaaca | 2 |
| MR934_prpC_Ver_F | Gacaacaactctcgaccctacaaatgataacaatgacgaggacaacatgag | 2 |
| Ec332_aceE_Ver_R | cgttccggatatcggcagcgcgacgaagtTgaagtgaccgaaatcctggtga | 2 |
| HSM25_pAJM.712-CML_marion_fwd | ctcggtaccaaattccagaaaagaggcctc | 3 |
| HSM26_pAJM.712_marion_rvs | Ctagtatttcccctctttctctagtattaa | 3 |
| HSM27_CML_marion_fwd | ttaatactagagaaagaggggaaatactagTAATAGAAATAATTTTGTTTAACTTTaa agttaagaggcaagaatggctcgtacctttaa | 3 |
| HSM28_CML_marion_rvs | Gaggcctcttttctggaatttggtaccgagttagaaatcgaactcaatac | 3 |
| HSM-45_SdhA_Mar_fwd | attttgtttaatactagagaaagaggggaaatactagTAATAGAAATAATTTTGTTTAAC TTTAAAGTTAAGAGGCAAGAatgggcagcagccatcacca | 3 |
| HSM-46_SdhB_Mar_rvs | cccctttcgggaggcctcttttctggaatttggtaccgagttacgcattacgttgcaaca | 3 |
| UGI_PaeI_R | gcttGCATGCttatagcatcttgatcttgttctctc | 3 |
| P02_pZ_R | ggcggatttgtcctactcag | 3 |
| P08_C2_v_F | gtggtattcactccagagcg | 3 |
| Ec541_sdhAint_fwd | gttgtgattggtgccggtgg | 3 |
| Ec660_pSEVA_rev | ATGGCTCATAACACCCCTTG | 3 |

| | | |
|---|---|---|
| HSM-48_FumA_Mar_rvs | cccctttcgggaggcctcttttctggaatttggtaccgagttatttcacacagcgggtgc | 3 |
| Ec1069_fumA_v_f | aaagtcgtactagtctcag | 3 |
| Ec1070_fumA_v_r | actgtattcatgacctgctc | 3 |
